# Microplastics are present in women’s and cows’ follicular fluid and polystyrene microplastics compromise bovine oocyte function *in vitro*

**DOI:** 10.1101/2022.11.04.514813

**Authors:** Nicole Grechi, Roksan Franko, Roshini Rajaraman, Jan B. Stöckl, Tom Trapphoff, Stefan Dieterle, Thomas Fröhlich, Michael J. Noonan, Marcia de A. M. M. Ferraz

**Author notes:** Corresponding author: Marcia de A. M. M. Ferraz, **Email:**.

## Abstract

The past several decades have seen alarming declines in the reproductive health of humans, animals and plants. While humans have introduced numerous pollutants that can impair reproductive systems (such as well-documented endocrine disruptors), the potential for microplastics (MPs) to be contributing to the widespread declines in fertility is particularly noteworthy. Over the same timespan that declines in fertility began to be documented, there has been a correlated shift towards a “throw-away society” that is characterised by the excessive consumption of single-use plastic products and a concomitant accumulation of MPs pollution. Studies are showing that MPs can impair fertility, but data have been limited to rodents that were force-fed hundreds of thousands of times more plastics than they would be exposed in the environment. As a first step to link *in vitro* health effects with *in vivo* environmental exposure, we quantified microplastics in the follicular fluid of women and domestic cows. We found that the concentrations of polystyrene microplastics that naturally occurred in follicular fluid were sufficient to compromise the maturation of bovine oocytes *in vitro*. Collectively, these findings demonstrate that microplastics may also be contributing to the widespread declines in fertility that have been occurring over recent Anthropocene decades.

## Introduction

Decades of careless use and improper disposal have resulted in plastic pollution accumulating almost everywhere on earth (Cole et al., 2011; Rochman and Hoellein, 2020), yet the health implications of the billions of tons of plastic pollution scattered across the planet’s terrestrial ecosystems are largely unknown. Microplastics (MPs) – defined as plastic particles <5 mm in size that are insoluble in water – have now been documented in even the most remote environments (Bergmann et al., 2019; Free et al., 2014; Lusher et al., 2015). The exponentially increasing volume of plastic pollution making its way into rivers, lakes and oceans has drawn considerable scientific, public, and governmental attention. However, despite the tens of thousands of peer reviewed publications on plastic pollution, an overwhelming majority of these have focused on aquatic ecosystems, and very little is known about the effects of MPs in terrestrial systems (Rillig and Lehmann, 2020). This knowledge gap is made all the more noteworthy by the fact that 80% of the Earth’s species live on land (Grosberg et al., 2012) and terrestrial systems may represent a larger environmental reservoir of MPs than oceans (de Souza Machado et al., 2018; Hurley and Nizzetto, 2018). With the plastic pollution crisis only expected to worsen over the coming decades (Kaza et al., 2018), it is imperative that we improve our understanding of the potential health effects of the tens of billions of tons of plastic pollution that litter the globe.

While any health effects are undesirable, the potential impacts of MPs on reproductive systems are of particular concern. Reproduction is central to the capacity for species to maintain stable populations. Poor reproductive health not only reduces individual fecundity but, if widespread, species survival. Over the past several decades there has been an alarming increase in the rates of reproductive dysfunctions and gamete abnormalities, reductions in gamete production, and altered embryo development in different species (Choi et al., 2016; D’Angelo and Meccariello, 2021; Du et al., 2016; Erdemir et al., 2014; Kumar et al., 2014; Rattan et al., 2017; Sone et al., 2004; Stuppia et al., 2015; Sussarellu et al., 2016a). While the overarching cause of these worrying declines in fertility has yet to be identified, emerging studies are showing that MPs represent a potentially serious threat to the reproductive health of terrestrial species (de Souza Machado et al., 2018; Hou et al., 2021; Ijaz et al., 2021; Jin et al., 2021). Findings have been limited to studies on laboratory animals, however, and the extent to which they are representative of the conditions that humans and animals are actually experiencing in the environment they inhabit is unknown (Mills et al., 2022). One of the primary reasons why existing *in vivo* MP studies are so far removed from reality is the lack of data on the bioaccumulation of MPs. At present there is only scattered information on how much plastic humans and other mammals ingest, and on how much is accumulating in different tissues and organs. This means that researchers interested in studying the health effects of MPs have no baseline exposure levels available to help guide their study design. Here, we assessed the extent to which MPs might be bio-accumulating in the follicular fluid of women and domestic cows. Particles isolated from human and bovine follicular fluid were identified via confocal Raman spectroscopy. We then investigated the effects of commercially available polystyrene MPs, on bovine oocyte *in vitro* at concentrations within the detection range of follicular fluid.

## Results and Discussion

### MPs, pigments, and plasticizers are present in human and cow follicular fluid

To perform a bio-mimetic experiment investigating the impact of MPs on fertility (and health in general) *in vitro*, it is necessary to have baseline information about what type, size, and amount of MPs can plausibly bio-accumulate in humans and other mammals. In reality, no such information is currently available. This can be attributed to two main reasons: 1) the vast majority of MPs research has focused on aquatic ecosystems, and 2) due to cost, time, and technological limitations there are no reliable methods for isolating, characterizing, and quantifying small MPs (<10 *μ*m) from complex biological samples. In an attempt to overcome the latter limitation, we optimized a protocol that allowed for the analysis of MPs in bovine and human follicular fluid (bFF and hFF, respectively). Nevertheless, there was still a large amount of undigested material in the samples (ranging from 4,912 total particles in water control 2 to 40,683 total particles in bFF3; Fig. 1a). Despite these large numbers of total detected particles, only about 5% of them had Raman spectra with hit quality index (HQI) ≥ 75.

**Figure 1.**
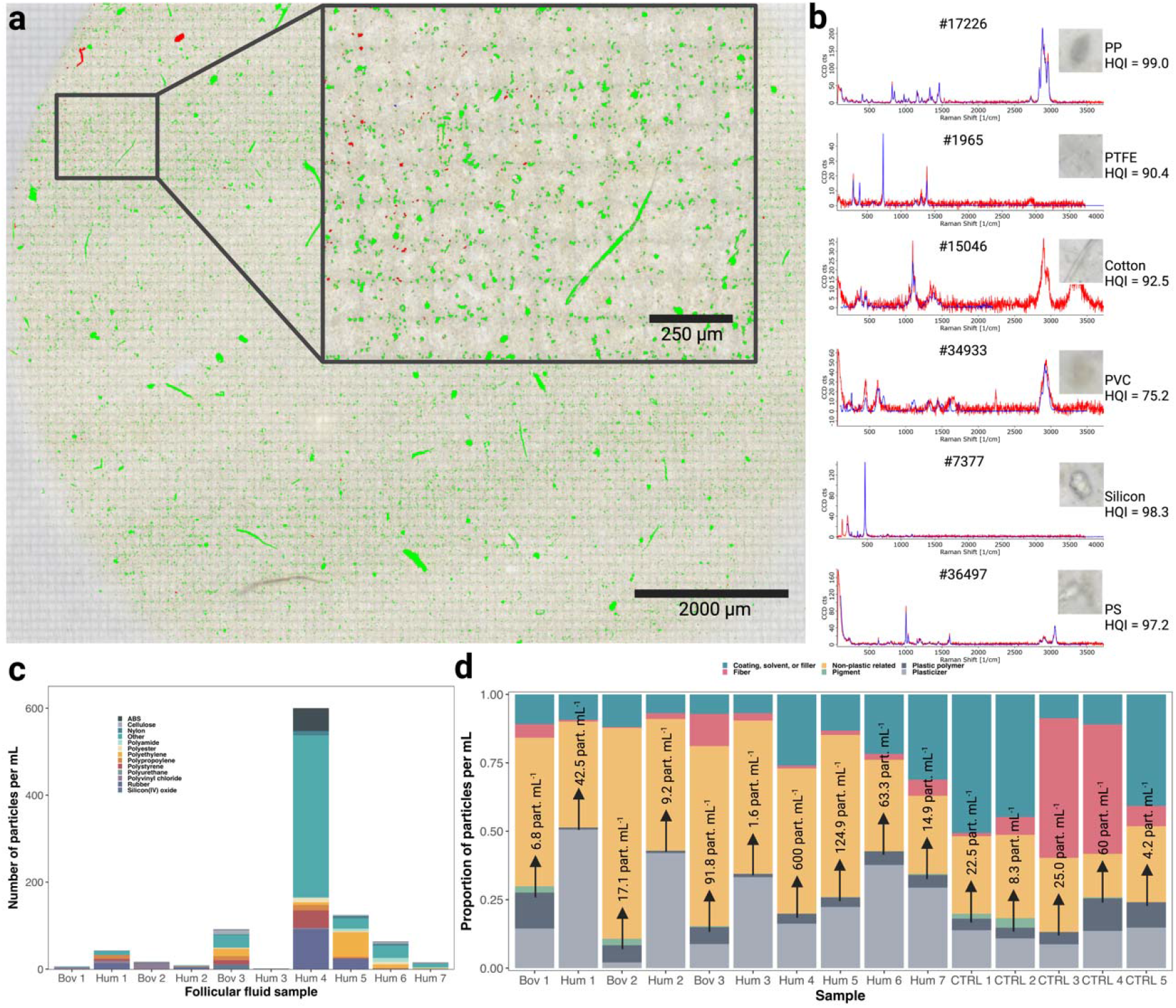
MPs were detected in bovine follicular fluid. In (**a**), an example of an imaged membrane (bFF3) that was used in the software Particle Scout to determine the number of particles and their sizes and for subsequent spectra match using the software TrueMatch is shown. Particles shown in green represent all counted particles, while those shown in red represent particles for which the spectra match had a hit quality index (HQI)≥ 75. In (**b**) examples of particles from bFF3 with their Raman spectra (red) and the matched Raman spectra of the corresponding plastic polymer (blue) from the ST Japan and/or SLoPP database. Particle number, polymer name and HQI are shown. In (**c**) the composition of MPs detected in the follicular fluid of humans (Hum) and bovine (Bov) by confocal Raman spectroscopy are shown. In (**d**) the counts of the different plastic and non-plastic analysed particles present in each sample are shown; written are the concentration of plastic polymers for each group and water controls. The number of MPs in FF in (**c** and **d**) was adjusted to the number of particles detected in their respective water control. PP = polypropylene, PTFE = polytetrafluoroethylene, PVC = polyvinylchloride, PS = polystyrene.

Although a large amount of undigested matter remained in the samples, an automated analysis of the sample membranes found MPs in all hFF and bFF samples, with mean concentrations of 122.3 and 38.6 MP particles mL^-1^, for humans and bovine respectively. Figure 1b shows examples of plastic polymers identified in FF and their spectra match. The total concentration of MP particles varied substantially between samples, with a total of 42.5 MP particles mL^-1^ in hFF1, 9.2 MP particles mL^-1^ in hFF2, 1.6 MP particles mL^-1^ in hFF3, 600 MP particles mL^-1^ in hFF4, 124.9 MP particles mL^-1^ in hFF5, 63.3 MP particles mL^-1^ in hFF6, 14.9 MP particles mL^-1^ in hFF7, 6.8 MP particles mL^-1^ in bFF1, 17.1 MP particles mL^-1^ in bFF2 and 91.8 MP particles mL^-1^ in bFF3 (Fig. 1c; Supplementary Table 1). Non-plastic related particles were the majority of identified particles (27.8 to 75.3 %), while MP polymers accounted for 0.8 to 23.5 % of the total number of identified particles. Other plastic-related particles that were identified included pigments (0 to 3.6 %), plasticizers (2.0 to 50.9 %), and coating, solubilizers and fillers (2.8 to 33.3 %; Fig. 1d). A total of 47 different MP polymers were identified, with rubber being the most abundant in humans, versus polyvinyl chloride in bovine. The average sizes of MPs in hFF was 11.5±12.2 μm in length (mean±SD) and 6.36±5.1 in width. In bFF, it was 11.4±7.2 μm in length and 6.3±5.1 μm in width. In all samples, most of the MPs particles were smaller than 20 μm (Supplementary Fig. 1 shows the size distribution of MPs in all samples).

**Table 1.** Differentially abundant proteins in oocytes incubated with or without polystyrene MPs. Protein name and ID, gene name, fold change and adjusted P-value are shown for oocytes incubated with 0.3 μm polystyrene beads compared to control and oocytes incubated with 1.1 μm polystyrene beads compared to the control. The full list of identified differentially abundant proteins is available in Data S2.

| Comparison | Protein name and ID | Gene name | Fold change | Adj. p-value |
| --- | --- | --- | --- | --- |
| 0.3 μm vs control | Major vault protein (Q3SYU9) | MVP | 0.6 | 8.2E-12 |
|  | Malate dehydrogenase, mitochondrial (Q32LG3) | MDH2 | 1.8 | 2.3E-07 |
|  | Acyl-CoA-binding protein (P07107) | DBI | 5.2 | 2.3E-05 |
|  | Protein-arginine deiminase (F1N404) | PADI6 | 0.7 | 4.6E-04 |
|  | Adenylate kinase 2 (P08166) | AK2 | 2.0 | 1.5E-02 |
|  | 10 kDa heat shock protein, mitochondrial (P61603) | HSPE1 | 1.5 | 1.5E-02 |
|  | NACHT, LRR and PYD domains-containing protein 5 (Q647I9) | NLRP5 | 0.8 | 2.3E-02 |
|  | Glutathione peroxidase 1 (P00435) | GPX1 | 1.9 | 2.3E-02 |
|  | Alpha-enolase (Q9XSJ4) | ENO1 | 1.2 | 3.2E-02 |
|  | Small nuclear ribonucleoprotein Sm D2 (Q3SZF8) | SNRPD2 | 2.6 | 3.6E-02 |
|  | H2A clustered histone 16, 7, 14, 10, 17 (A0A3Q1N7K2) | H2AC16; H2AC7; H2AC14; H2AC10; H2AC17 | 1.5 | 4.2E-02 |
|  | Triosephosphate isomerase (Q5E956) | TPI1 | 1.4 | 4.5E-02 |
|  | Thioredoxin (O97680) | TXN | 2.0 | 4.7E-02 |
| 1.1 μm vs control | ATPase inhibitor, mitochondrial (P01096) | ATP5IF1 | 3.5 | 1.8E-08 |
|  | Protein-arginine deiminase (F1N404), | PADI6 | 0.7 | 2.1E-04 |
|  | Major vault protein (Q3SYU9) | MVP | 0.8 | 4.7E-04 |
|  | Heteroous nuclear ribonucleoprotein (A0A3Q1MJD5) | HNRNPM | 1.5 | 4.7E-04 |
|  | Tubulin-specific chaperone A (P48427) | TBCA | 2.4 | 1.5E-03 |
|  | Desmoglein-1 (Q03763) | DSG1 | 18.8 | 3.0E-03 |
|  | Tropomyosin alpha-1 chain (Q5KR49) | TPM1 | 1.9 | 3.9E-03 |
|  | Splicing factor proline and glutamine rich (E1BQ37) | SFPQ | 1.6 | 1.6E-02 |
|  | Heat shock protein HSP 90-alpha (Q76LV2) | HSP90AA1 | 0.7 | 2.3E-02 |
|  | Probable ATP-dependent RNA helicase (F1MBQ8) | DDX5 | 2.8 | 2.4E-02 |
|  | Desmoplakin (E1BKT9) | DSP | 10.9 | 2.4E-02 |
|  | BCL2 associated transcription factor 1 (F1MQU4) | BCLAF1 | 2.6 | 3.9E-02 |
|  | NACHT, LRR and PYD domains-containing protein 5 (Q647I9) | NLRP5 | 0.8 | 3.9E-02 |

Based on their sizes and densities, we determined that MPs summed to 0.0309 μg mL^-1^ in hFF1, 0.0032 μg mL^-1^ in hFF2, 0.0002 μg mL^-1^ in hFF3, 0.3869 μg mL^-1^ in hFF4, 0.026 μg mL^-1^ in hFF5, 0.042 μg mL^-1^ in hFF6, 0.088 μg mL^-1^ in hFF7, 0.0009 μg mL^-1^ in bFF1, 0.005 μg mL^-1^ in bFF2 and 0.069 μg mL^-1^ in bFF3. On average, we detected 122.3 particles mL^-1^ or 0.102 *μ*g mL^-1^ in hFF and 38.6 particles mL^-1^ or 0.025 *μ*g mL^-1^in bFF.

In general, 7 polymers were the most abundant in human FF (polyvinyl chloride – PVC, polyethylene – PE, polystyrene – PS, polypropylene – PP, polyurethane – PU, rubber – RUB, and ABS). While in bovine FF 10 polymers (PVC, PE, cellulose acetate– CE, PS, PP, silicon – SL, nylon – NL, polyester – PES, ABS and polyamide – PA) were the most abundant. The concentrations of MPs we found in human and bovine FF were greater than previously detected in human placentae (0.52 particles g^-1^; PE, PP and Polyurethane – PU), human lungs (0.69 particles g^-1^; PP, polyethylene terephthalate – PET, Polytetrafluoroethylene – PTFE, PS, polyacrylonitrile – PAN, polymethyl methacrylate – PMMA and PU), and raw cow milk (4.1 particles mL^-1^; PE, PES, PP, PTFE, PU and PA) (Braun et al., 2021; Da Costa Filho et al., 2021; Jenner et al., 2022; Ragusa et al., 2021) via particle spectroscopy, but less than in human blood (1.6 *μ*g mL^-1^; PET, PE, PE and PMMA), cow raw milk (∼18.1 *μ*g mL^-1^; PVC, PP, PE, PS and PMMA), cow blood (∼5.4 *μ*g mL^-1^, PVC, PP, PE, PS and PMMA), and cow meat (∼420.5 *μ*g mL^-1^, PVC, PP, PE, PS and PMMA) via pyrolysis – gas-chromatography/mass-spectroscopy (Py-GC/MS) (Leslie et al., 2022; Veen et al., 2022). Such differences are likely due to the fact that, by using spectroscopy in our study, we were limited by the lower 0.3 *μ*m detection limit of Raman spectroscopy, and were likely under-estimating the true number of MPs contained in the samples. While the analysis of MPs by Py-GC/MS can detect particles down to monomers, which is more sensitive to the amount of plastics, it comes with the trade-off of not being able to determine the size and shapes of particles. This latter information can be critical for understanding the potential for MPs to cross biological barriers, and why we opted to use Raman spectroscopy in our analyses.

Of importance for the present study, from the total 0.102 *μ*g mL^-1^ and 0.025 *μ*g mL^-1^ of MPs we detected in human and bovine FF, 0.0013 and 0.0043 *μ*g mL^-1^, or 1.4 and 17.3%, were PS, respectively. The percentage of MPs we identified as PS in bovine FF was similar to that previously detected in cow blood (13%; 0.7 *μ*g g^-1^), though previous reports on MPs in cow meat and milk found a lower proportion of PS (5.11%; 21.5 *μ*g g^-1^, and 0.11%; 0.02 *μ*g g^-1^ respectively) (Veen et al., 2022). In a study on human blood, PS was the primary polymer detected (60%; 0.96 *μ*g mL^-1^) (Leslie et al., 2022). Full details on the composition of each of the samples are shown in Data S1, and MP polymers are summarized in Supplementary Table 1.

It is important to highlight, as seen for human and bovine blood, bovine milk and meat, the total amount of MPs detected is extremely variable between individuals and animal origin. Here, we have shown that, similar to other bodily fluids and tissues, MPs are also present in human and bovine follicular fluid. Nonetheless, it is still unknown to what extent the concentrations of MPs in FF might correlate with the amount of MPs in less invasive blood and/or stool samples, nor whether these inter-individual differences are related to other factors such as diet, or age, which would be an important topic of future research.

It is also important to highlight, that the isolation and characterization protocol used for the detection of MP particles in FF, required 3-7 days per sample to acquire the Raman spectra of all the detected particles, in addition to 4-18 hours to match these to the database. While this protocol was sufficient for the scale of the present study, it is clearly not a viable option for large scale applications. Showing that the refinement of methodologies for the sampling, detection, and quantification of MPs from biological samples is critical for the future of the field

### Polystyrene MPs reduce oocyte maturation and induce zona pellucidae damage

While the present work is not the first study on the effects of MPs on mammalian reproduction, it is the first to demonstrate the effects of MPs at concentrations that fall within the weight of MPs detected in reproductive fluids. For instance, in a study where mice were fed with 15 – 1,500 μg of polystyrene day^-1^ for 90 days, they exhibited a reduced ovarian follicle reserve, ovarian oxidative stress, granulosa cell apoptosis, and ovarian fibrosis (An et al., 2021). While the results of such studies are certainly alarming, the extent to which they are representative of the conditions humans and animals are actually experiencing in the real world is questionable. More specifically, a recent study where mice were fed 30 mg kg^-1^ of body weight of PS MPs for 35 days, the concentration of PS detected in their blood (135.86 *μ*g mL^-1^), was 141 and 194 times higher than the concentration of PS MPs found in human and cow blood, respectively (Liu et al., 2022). Those authors found that MPs of ∼0.8 μm can accumulate in ovaries and cause increased oxidative stress and reduced oocyte maturation (Liu et al., 2022), but they detected PS MPs in mouse ovarian tissue at a concentration of 62.6 *μ*g g^-1^, which was 48,153 and 14,558 times higher than the amount of PS MPs we identified in human and bovine FF, and 613 and 2,504 times higher than the total concentration of MPs, respectively. Using values in the range of MPs weight detected in bovine FF and human FF, we demonstrated that the negative effects of MPs on oocytes can actually be seen at an exposure to lower amounts of MPs, and within a much shorter, 24 hour, incubation time.

Here, we used polystyrene beads of 0.3 (0.01 μg mL^-1^) and 1.1 μm (0.1 μg mL^-1^), since these were the most abundant MPs in human blood (Leslie et al., 2022). The oocyte *in vitro* maturation performed in the presence of polystyrene beads resulted in a significantly lower number of mature oocytes in comparison to the control group (control: 68.1±10.3%, 0.3 μm MPs: 42.0±14.8%, and 1.1 μm MPs: 41.0±17.6%; p < 0.00001 for both 0.3 and 1.1 μm MPs; Fig. 2a), with no significant difference between the 0.3 and 1.1 μm MP treatments (p = 0.963). The number of degenerated oocytes varied between 27% and 37% and did not differ between any of the groups, however the control group had significantly fewer oocytes with broken zona pellucidae, even though they were submitted to same handling as the COCs incubated with MPs, when compared to the groups incubated with MPs (control: 4.4±5.2%, 0.3 μm MPs: 21.4±7.6%, and 1.1 μm MPs: 26.2 ±10.9%; p < 0.0001 for all groups). Notably, of the oocytes with broken zona pellucidae, 80% were mature in the control group, as compared to 40 and 35%, for the 0.3 and 1.1 μm MP treatments, respectively, but these oocytes were categorised as having broken zona pellucidae, and not as mature.

**Figure 2.**
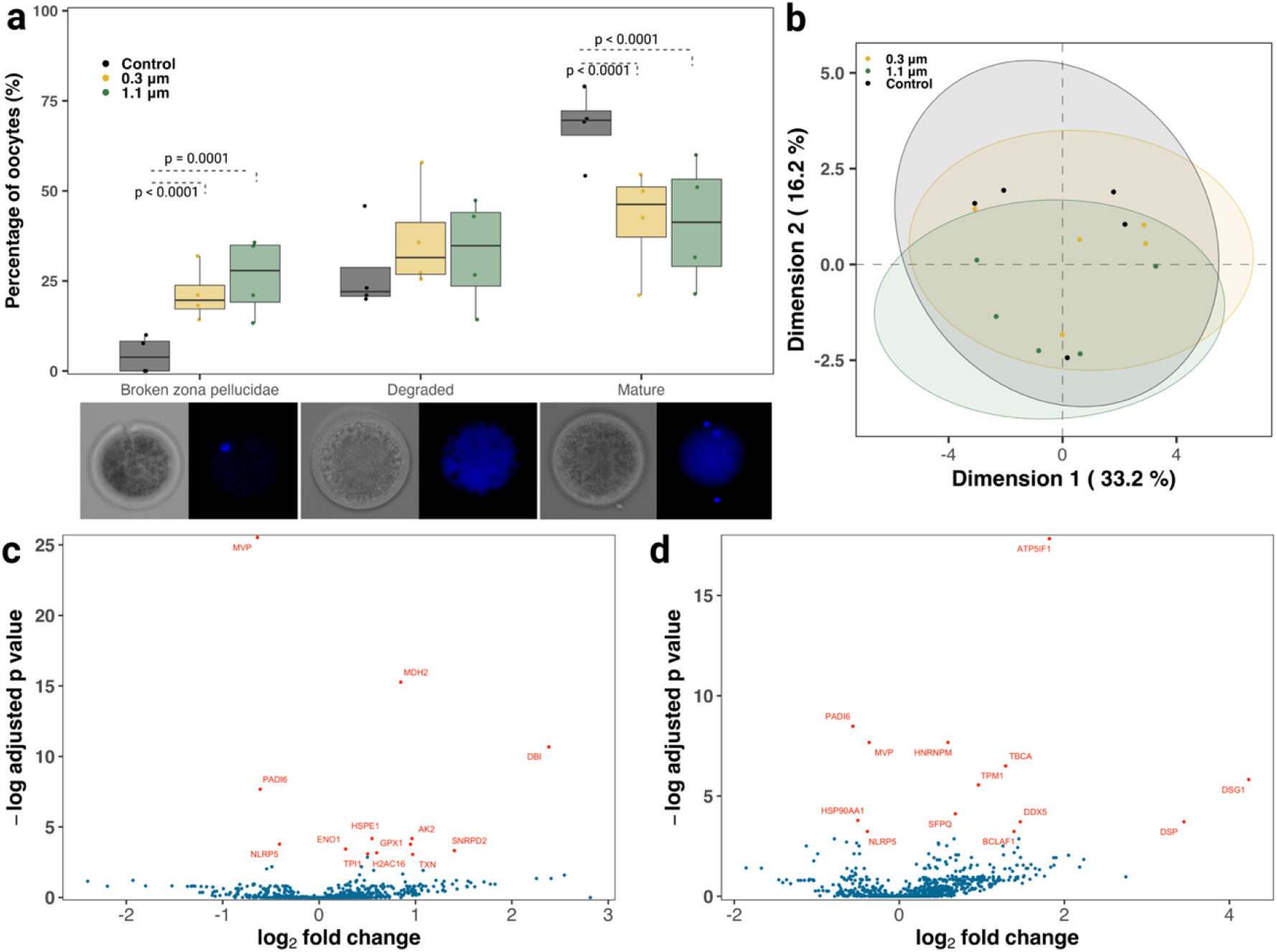
Polystyrene MPs negatively influenced oocyte maturation. The boxplots in (a) show the effects of 24 h exposure to 0.3 and 1.1 μm polystyrene beads on oocyte maturation *in vitro*. Image examples of each stage characterization below the panel, oocytes were stained for DNA (Hoechst33342, blue). The scatter plot in (b) depicts the first two dimensions of a principal component analysis (PCA) across the proximity matrix of a random forest model classifying oocytes exposed to 0.3 or 1.1 μm polystyrene beads versus the control, based on protein abundance profiles. Volcano plots depicting differently abundant proteins between oocytes incubated for 24 h in the presence of (c) 0.3 μm and (d) 1.1 μm polystyrene beads compared to a control are also shown.

### PS MPs alter the oocyte proteome

Beyond these functional responses, proteomics analysis of oocytes incubated with or without microplastic also revealed proteins altered in abundance that are part of critical oocyte function pathways. We identified 2,060 proteins across all bovine oocyte samples and a low variance was observed between the different groups (Fig. 2b, Supplementary Figure 2). A total of 13 proteins were present at significantly different abundances when comparing oocytes incubated with 0.3 μm PS beads to the control group (3 down-regulated and 10 up-regulated), while 13 proteins differed significantly between oocytes incubated with 1.1 μm PS beads as compared to the control (4 down-regulated and 9 up-regulated; Fig. 2c and d; Table 1).

Proteins that are known to act upon oxidative stress were up-regulated in the presence of MPs: TXN (Jiang et al., 2020), GPX1 (Chu et al., 2020), ATP5IF1 (García-Aguilar and Cuezva, 2018) and PADI6 (Kan et al., 2011; Yurttas et al., 2008), which together with the up-regulation of SNRPD2 (Wyatt et al., 2022), known for promoting DNA repair response, indicate that the presence of MPs may induce oxidative stress and DNA damage in incubated oocytes during *in vitro* maturation. Proteins known to induce apoptosis, such as BCLAF1 (Kasof et al., 1999), TPM1 (Tang et al., 2018) and AK2 (Lee et al., 2007) were up-regulated in MPs incubated oocytes, while the MVP (Sutovsky, 2009), a known oocyte protector, was down-regulated. Additionally, DDX5 (Lan et al., 2022) was up-regulated and HSP90AA1 (Liu et al., 2018) was down-regulated in the presence of MPs. Together with an up-regulation of HNRNPM (Chen et al., 2017; Kogasaka et al., 2013) and TBCA (Nolasco et al., 2005), and down-regulation of PADI6 (Kan et al., 2011; Yurttas et al., 2008), these differentially abundant proteins can provide a mechanistic explanation as to the reduced maturation rate observed in oocytes exposed to MPs, since these proteins are important for nuclear maturation, spindle formation and microtubule and organelle organization (summarized in Fig. 3). Nevertheless, the low number of proteins differentially expressed between the 2 groups, indicates that the cumulus cells must play an important role on controlling the effects of PS beads on oocyte, and it should be further investigated.

**Figure 3.**
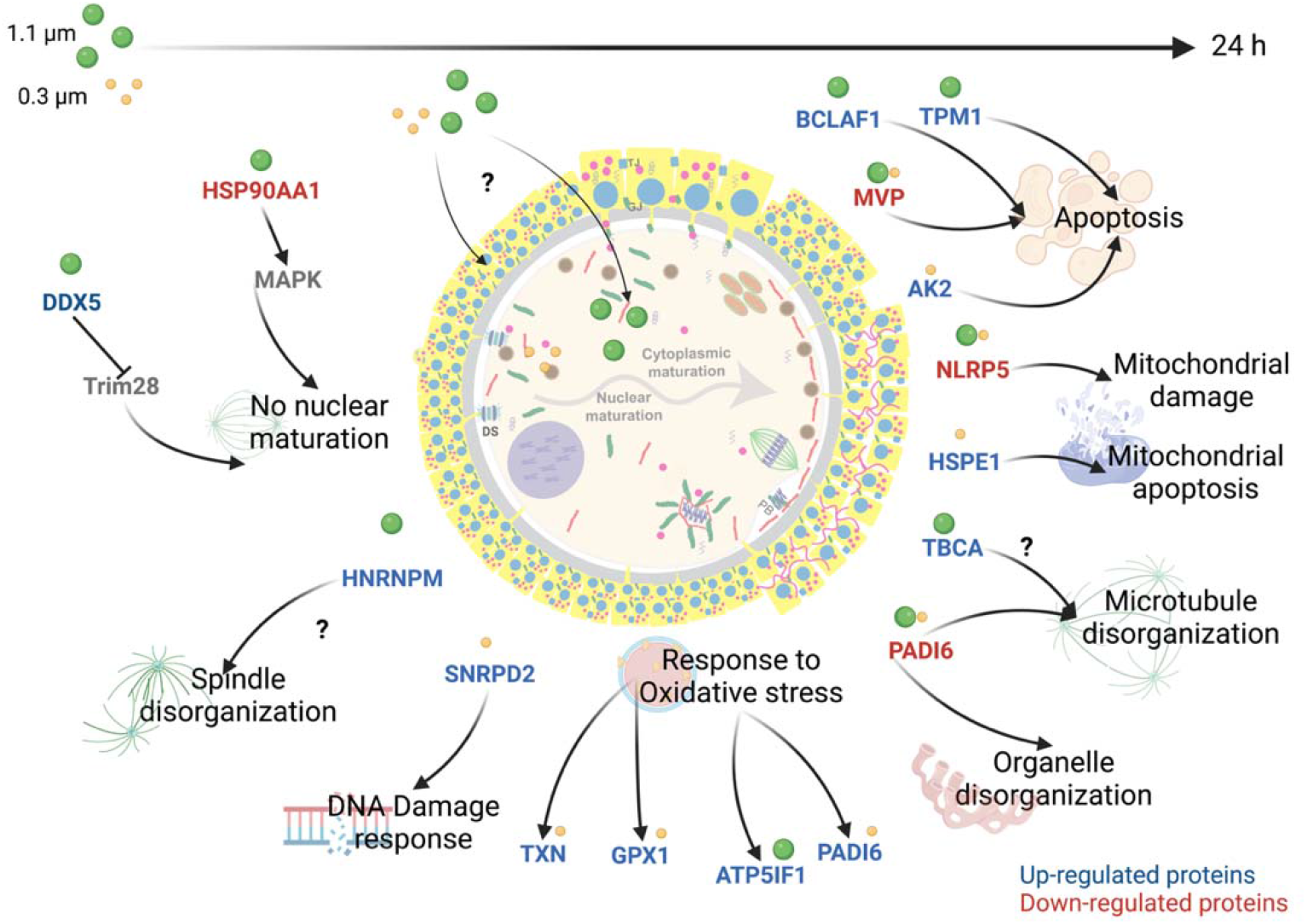
Summary of the effects of polystyrene MP on oocyte proteomes. Differentially expressed proteins in oocytes incubated for 24 h with 0.3 (yellow) and 1.1 (green) μm polystyrene beads compared to control are shown. Proteins were found to regulate major pathways in oocyte function, such as apoptosis, mitochondrial damage, mitochondrial apoptosis, microtubule disorganization, organelle disorganization, spindle disorganization and nuclear maturation. Moreover, proteins part of response to oxidative stress and DNA damage were also differently expressed. Figure made using BioRender.

## Conclusions

The ubiquitous and long-lived nature of MPs has made them synonymous with the seemingly irreversible mark of mankind on our planet. While evidence is still extremely limited, emerging studies are showing that MPs represent a potentially serious threat to the reproductive health of laboratory rodents and aquatic species (An et al., 2021; de Souza Machado et al., 2018; Deng et al., 2021; Hou et al., 2021; Ijaz et al., 2021; Jaikumar et al., 2019; Jeong et al., 2016; Jin et al., 2021; Li et al., 2021; Sussarellu et al., 2016b). Due to the lack of a reliable method for isolating and characterising MPs from biological tissues and fluids, however, our capacity to investigate the presence of MPs in naturally occurring biological samples has been limited. In addition, there were heretofore no studies linking MP contamination to changes in fertility in non-lab-rodent terrestrial mammals. This study provides the first comprehensive evaluation of the detrimental effects of MPs contamination on the female reproductive system. Using a protocol that was optimized for the isolation of small MPs from FF, we have shown that MPs are present in the FF of women and domestic cows. From our *in vitro* analyses, we found that polystyrene, in the concentrations of MPs detected in FF were shown to compromise the normal functioning of female bovine gametes. Collectively, these findings provide evidence as to how the billions of tons of MP pollution may be contributing to the widespread increase in the rates of reproductive dysfunctions that have been increasing over recent decades (Choi et al., 2016; D’Angelo and Meccariello, 2021; Du et al., 2016; Erdemir et al., 2014; Kumar et al., 2014; Rattan et al., 2017; Sone et al., 2004; Stuppia et al., 2015; Sussarellu et al., 2016a). Crucially, and addressed here, the discrepancy between the concentrations of MPs used in existing studies versus those that humans and animals can experience in the environment, emphasizes the dire need to better understand how and where terrestrial animals and humans bioaccumulate MPs. This information would provide a more realistic picture of the impact of MP pollution on terrestrial ecosystems, and allow researchers to establish baseline levels of exposure for *in vitro* experiments on the impact of MPs on health and fertility. Humans and animals have been exposed to MPs for decades. The short-term effects of polystyrene MPs on oocytes *in vitro* we observed here are, therefore, likely to be more pronounced *in vivo*, due to the detrimental effects of MPs compounding over time. Future investigations should evaluate the effects of different plastic polymers on the oocyte and how the different amounts/types of MPs in FF can influence IVF outcomes.

## Materials and Methods

### MP contamination prevention

To prevent sample contamination with airborne/materials MPs, all procedures were performed in a laminar flow hood, and all flasks and other apparatuses were replaced by glass materials whenever possible. Moreover, all materials and equipment used were rinsed three times with filtered (0.1 μm filter – Merck Isopore) ultra-pure water prior to use. All reagents and water used in the protocols described below were also filtered using a 0.1 μm filter before use. Samples/vials were closed all the time using aluminium foil. All materials used for human and bovine follicular fluid aspiration were new and sterile (by the manufacturer) and procedure blanks using such materials were performed, as described in the microplastic isolation section below.

### Microplastic polystyrene beads

Polystyrene beads (SURF-CAL™ particle size standards) having average sizes of 0.047 μm, 0.100 μm, 0.304 μm and 1.112 μm were purchased from Thermo Fisher Scientific. The beads were obtained in deionized filtered water in a concentration of 3 × 10^8^ particles mL^-1^ and were diluted according to the weight required for each experiment.

### Human follicular fluid collection

Follicular fluid samples (n=7) were collected from patients undergoing intracytoplasmic sperm injection (ICSI) treatment between January and September 2022, at the Dortmund Fertility Centre, Germany. The study was approved by the ethics committee at Witten Herdecke University (no. S-262/2021). Briefly, aspiration of cumulus oocytes complexes (COCs) was done ultrasound-guided from large Graafian follicles after ovarian stimulation and hCG priming. The oocytes were processed for further ICSI-treatment. The extant hFF without cumulus oocytes complexes was transferred to pre-washed and sterile glass bottles (Schuett biotec, Germany) and immediately frozen at −20°C. hFF collection and processing were performed under clean room facilities.

### Bovine oocyte and follicular fluid isolation

Bovine ovaries were obtained from a local slaughterhouse and immediately transported to the laboratory in a stainless-steel container at room temperature. Immature cumulus oocytes complexes (COCs) were aspirated from follicles with a size between 2 and 8 mm, together with follicular fluid, by using a vacuum pump and a 19G needle. Collected fluids with COCs from ∼20-30 ovaries (N = 3 pools) were let to pellet for a maximum of 10 min. The pellet was then transferred to a petri dish with the equivalent amount of washing media (IVF Bioscience, UK) for oocyte selection, and remaining follicular fluid frozen in the pre-washed glass vials at −20°C for MPs isolation as described below. Each bFF pool was collected in a different day, each pool comprises animals coming from the same farm, and pools from different days include animals from different farms.

### Microplastic isolation from human and bovine follicular fluid

Three pools of bovine follicular fluid (bFF) and follicular fluid from seven human (hFF) patients, left after removal of the COCs, were used for microplastic isolation. Water controls from follicular fluid aspiration (N=2 for bFF and N=3 for hFF) were produced by, aspirating filtered ultra-pure water in the same systems that the human and bovine follicular fluids were aspirated, the same day the samples were processed for digestion. On day one, 2 mL of each sample were added to an Erlenmeyer for digestion in KOH 10% in a proportion of 1:25 (sample:digestion solution). The samples were incubated in a shaker at 60°C and 250 rpm for 24 h. After this period, NaClO was added to each digestion to reach a final concentration of 0.84%, and samples were incubated for another 24 h in the shaker at 60°C and 250 rpm. On day three, all samples were filtered in a 47 mm polytetrafluoroethylene polymer (PTFE) membrane (0.45 μm pores, Merck Millipore, USA) and rinsed with filtered ultra-pure water at least three times in order to prevent any NaClO and KOH contamination. Next, the membranes were placed in a beaker containing 50 mL of HNO_3_ 20% and incubated in an ultrasonic bath (TI-H-5 MF2 230 V, Elma Schmidbauer GmbH, Germany) with 100% power, sweep function and a frequency of 45 kHz for 15 min, to transfer plastics/undigested matters from the membrane into the solution. The resulting solution was then kept in the shaker at 40°C and 250 rpm for another 24 h. At the end of the third digestion day, the samples were filtered using 13 mm PTFE membranes (0.45 μm pores) and again rinsed three times with ultra-pure filtered water. The membranes were then mounted on a glass slide for microscopy and spectroscopy analysis.

The blank control waters were processed using same protocol and in parallel to the corresponding follicular fluids, including its aspiration using same materials/equipment: water control 1 with bFF1 and bFF2 samples, water control 2 with bFF3 sample, water control 3 with hFF1 sample, water control 4 with hFF2, hFF3 and hFF4 samples, and water control 5 with hFF5, hFF6 and hFF7 samples. Adjustment of MPs detection on FF samples to water controls (as described below) was performed using these corresponding water-FF samples combinations.

### Oocyte isolation, incubation with MPs and nuclear stage analysis

After collection as described above, the Petri dish with COCs was screened with a stereomicroscope and good quality oocytes (intact oocyte with homogeneous cytoplasm and, at least, 3 layers of cumulus cells) were selected. COCs were then washed three times in washing media, and one time in BO-IVM (IVF Bioscience, UK), before being randomly assigned to one of three incubation groups: 1) a control group containing only maturation media (N = 114); 2) media containing PS beads of size 0.304 μm (N = 93); and 3) media containing PS beads of size 1.112 μm (N = 103). The concentration of the beads in each group was 0.01 and 0.1 μg mL^-1^ for beads 0.3 and 1.1, respectively. The groups were cultured for 24 h in an incubator at 38.5°C in a humidified atmosphere of 5% CO_2_ and 95% O_2_. After the 24 h period of incubation, the oocytes were denuded by pipetting to remove cumulus cells, washed, and either fixed in paraformaldehyde 4% for nuclear staging (N = 3 replicates) or frozen at −80°C in pools of 9 oocytes for proteomics analysis (N = 5 pools per group, from 3 replicates).

For determining oocyte nuclear stage, fixed oocytes were washed in phosphate buffer saline (PBS), stained with Hoechst 33342 (5 μg mL^-1^) for 45 min and imaged in an EVOS M7000 Microscope using a ×40 NA 1.25 objective. The oocytes were analysed for nuclear stage (metaphase 1 or 2 – determined by the presence of an aligned metaphase plate with/without a polar body, respectively), degenerated (no visible nuclear material, or pyknotic nucleus), and broken zona pellucidae (visible breaks in the zona pellucida).

### Oocyte proteomics analysis

For lysis, nine oocytes per sample were sonicated in lysis buffer (8 M urea in 50 mM ammonium bicarbonate). Samples were then reduced in a final concentration of 4 mM dithiothreitol and 2 mM Tris(2-carboxyethyl)phosphine hydrochloride for 30 min at 56°C. Alkylation was done in the dark for 30 min with 8 mM iodoacetamide. Residual iodoacetamide was quenched by adding dithiothreitol to a final concentration of 10 mM. Samples were then digested in two steps, first 4 h with LysC at 37°C, then after dilution to 1 M Urea, trypsin was added and samples were digested overnight at 37°C. Prior to LC-MS/MS analysis, samples were dried using a centrifugal evaporator and then resuspended in 0.1 % formic acid (FA). Peptides were analysed with a Q Exactive HF-X (Thermo Fisher Scientific) mass spectrometer coupled to an Ultimate 3000 RSLC chromatography system (Thermo Fisher Scientific). First, samples were trapped (PepMap 100 C18, 100 μm × 2 cm, 5 μM particles; Thermo Fisher Scientific) at a flow of 5 µL min^-1^ of 0.1 % FA. Peptides were then separated (EASY-Spray; PepMap RSLC C18, 75 μm × 50 cm, 2 μm particles, Thermo Fisher Scientific) at a flow rate of 250 *η*L min^-1^ employing a two-step gradient: First ramping from 3 % B to 25 % B in 80 min then to 40 % B in 9 min (A: 0.1% FA in water, B: 0.1 % FA in acetonitrile). The mass spectrometer was run in data dependent acquisition mode with up to 15 MS/MS scans per survey scan. Raw files were processed with MaxQuant 1.6.7.0(Tyanova et al., 2016) using the bovine subset of both Swiss-Prot and TrEMBL (Retrieval date: 24/05/2022) as database. Additionally, the built-in common contaminants database was used. Label free quantification and match between runs was turned on. Subsequent statistical analysis was done using R (4.2.0). Differentially abundant proteins were identified with the package MS-EmpiRe (Ammar et al., 2019) as described in (Flenkenthaler et al., 2021).

### Confocal Raman spectroscopy of isolated microparticles and polystyrene beads

A confocal Raman (alpha300 R, WITec, Germany) with a spectrometer (UHTS 300, WITec, Germany), and a 532 nm laser was used for the characterization of the microplastics. The microscope was operated by using the Control Five software, and the acquired data were processed by the Project Five software for compensation of background noise or cosmic radiation signal. Initially, the microscope was focused on the samples by using a ×50 NA 0.75 Zeiss objective, and the whole area of the membrane was imaged using the ‘Area Stitching’ option and the Z-stack auto-focus. Then, the Particle Scout software was used to identify all the particles present in the imaged membrane. The Raman spectra of each identified particle was then measured with a ×100 DIC NA 0.9 Zeiss objective, using an autofocus function, with a laser power of 10 mV, an accumulation of 4 and integration time of 0.45 s. The presence of microplastics was confirmed by matching the spectra found using the True Match-Integrated Raman Spectra Database Management software, with the S.T. Japan database (S.T. Japan Europe GmbH, Germany) and the SloPP MPs Raman library (Munno et al., 2020). Only matches with a hit quality index (HQI) value higher than 75% were considered for further analysis. Poly(tetrafluoroethylene-co-perfluoro-(alkyl vinyl ether)) – PTFE – and Perfluoroalkoxy alkane – PFA – particles were excluded from the analysis, since they are components of the pore membrane used for imaging. After matching and excluding membrane particles, the number of MPs was adjusted to the particles detected in the water controls by subtracting the number of particles detected in the water controls from the number of particles in FF, for each specific polymer detected individually.

All identified particles were classified as: 1) non-plastic related particles; 2) plastic polymers; 3) plasticizers; 4) pigments; 5) coatings, solvents or fillers (CSF); 6) fiber; and 7) unknown. Data S3 show a list of all identified particles and their classification. Particle number was converted to weight by using the formula:

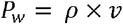

where P_w_ is particle weight, *ρ* is the particle density, and *v* is the particle volume. See Supplementary file Table 2 for full information on the density of the particles analysed here.

### Statistical analysis

We were interested in studying the effects of MPs on sperm and oocyte function, as well as on oocyte proteomics. Oocyte maturation data consisted of percentages that ranged between 0 and 100. We therefore assessed the effects of MPs on this functional trait via generalised linear regression models with a quasibinomial error distribution to account for the fact that the data were under-dispersed. In addition, we also applied a hierarchical approach in which data from each animal and replicate were allowed to have randomly varying intercepts. Significance for pair-wise combinations of discrete variables was checked using a Tukey HSD. Differences were considered significant when p < 0.05. All data analysis and visualization were carried out in R (ver. 4.2.1), and the scripts and packages used for carrying out our analyses are described in supplementary file S1 as well as on the GitHub repository https://github.com/NoonanM/MPs_and_Fertility.

Statistical analysis of the proteomics data was also performed using R. Fold changes and p-value calculations were performed with the package MS-EmpiRe (Ammar et al., 2019). Proteins abundances were considered significantly differentially if the adjusted p-value < 0.05. The differentially expressed proteins identified via these analyses are shown in Table 1. In addition, we used a random forest model to classify oocytes as belonging to either the control group, the 0.3 μm polystyrene bead treatment, or the 1.1 μm polystyrene bead treatment. The model was fit using the R package randomForest (Liaw and Wiener, 2018), using 20,000 trees, sampled with replacement, and five candidate genes were sampled at each split. We then evaluated the within-sample classification accuracy of the model and assessed the relative importance of the individual genes in overall model performance.

## Supporting information

R code used for all analysis

Full list of all particles detected in each sample

Proteomics data

Particles classification list

## Data availability

All data are available in the main text or the supplementary materials. The mass spectrometry proteomics data have been deposited to the ProteomeXchange Consortium via the PRIDE (Perez-Riverol et al., 2022) partner repository with the dataset identifier PXD036415, accessible via: https://www.ebi.ac.uk/pride/.

## Acknowledgments

We would like to thank Bachuki Shashikadze for his input in bioinformatics. This research was supported by LMUexcellent, funded by the Federal Ministry of Education and Research (BMBF) and the Free State of Bavaria under the Excellence Strategy of the Federal Government and the Lalnder. MAMMF was also supported by the Alexander von Humboldt Foundation in the framework of the Sofja Kovalevskaja Award endowed by the German Federal Ministry of Education and Research. MJN was supported by NSERC Discovery Grant RGPIN-2021-02758. TF was supported by the SFB 1357 funded by the Deutsche Forschungsgemeinschaft (DFG) Project number 391977956. TT and SD were supported by Theramex Germany GmbH.

## Author Contributions

Conceptualization: MAMMF; Methodology: MAMMF, NG, TF, JS, MJN and TT; Investigation: NG, RF, RR, JS; Supervision: MAMMF, MJN, TF, TT and SD; Writing— original draft: NG, MAMMF, MJN; Writing—review & editing: all authors.

