## Supplementary material for "Microplastics are present in women’s and cows’ follicular fluid and polystyrene microplastics compromise bovine oocyte function *in vitro*": R code used for all analysis

Supplementary File S1. R code used to generate the results and figures presented in the main text.


### Supplementary File S1. R code used to generate the results and figures presented in the main text.

###### Nicole Grechi, Roksan Franko, Roshini Rajaraman, Jan B. Stöckl, Tom Trapphoff, Stefan Dieterle, Thomas Fröhlich, Michael J. Noonan, Marcia de A. M. M. Ferraz

---

### Preface

This file details the statistical analyses used to assess the effects
of microplastics on oocyte function *in vitro*.

##### Packages used

The packages used in these analyses are detailed below. Package
versions are included at the end of this document.

```
# Packages used in the analyses
library(MASS)
library(lme4)
library(multcomp)
library(MuMIn)
library(gtsummary)
library(ggplot2)
library(knitr)
library(ggrepel)
library(VennDiagram)
library(FactoMineR)
library(ellipse)
library(randomForest)
library(caret)
library(ggridges)
library(gridExtra)
library(tidyverse)
library(msEmpiRe)
library(Biobase)
library(limma)
```

---

### Oocytes

The first step in the analyses was to characterise the microplastics
(MPs) in bovine follicular fluid, and assess the effects of MPs exposure
on oocytes.

#### MPs, pigments, and plasticizers in cow follicular fluid

We first wanted to visualise the different plastic polymers that were
identified in bovine follicular fluid samples, as well as in the water
controls. Each of these samples came from a pool of follicular fluid
collected from different cows, with each sample belonging to a different
collection period. For the figures presented in the main text, we
removed information on particles that were non-plastic related. Details
on the non-plastic particles are presented in Data S1.

```
# Import the plastic particle data
data <- read.csv("Data/Particle_categories.csv")
data2 <- read.csv("Data/MP_fluids.csv")
```

```
C <- 
  ggplot(data = data2, 
         aes(x = Sample,
             y = particles_ml,
             fill = Microplastic)) +
  geom_bar(stat = "identity", alpha = 0.8) +
  scale_fill_manual(labels = c("ABS",
                               "Cellulose",
                               "Nylon",
                               "Other",
                               "Polyamide",
                               "Polyester",
                               "Polyethylene",
                               "Polypropoylene",
                               "Polystyrene",
                               "Polyurethene",
                               "Polyvinyl chloride",
                               "Rubber",
                               "Silicon(IV) oxide"),
                    values = c("#001219", "#8d96a3", "#005F73", "#0A9396", "#94D2BD",
                               "#E9D8A6","#EE9B00", "#CA6702",  "#9B2226",  "#6d597a", "#355070",  "#fbb13c", "#bf0603")) +
  theme_bw()+
  theme(panel.grid.major = element_blank(),
        panel.grid.minor = element_blank(),
        axis.title.y = element_text(size=14, family = "sans", face = "bold"),
        axis.title.x = element_text(size=14, family = "sans", face = "bold"),
        axis.text.y = element_text(size=12, family = "sans"),
        axis.text.x  = element_text(size=12, family = "sans"),
        plot.title = element_text(hjust = -0.05, size = 10, family = "sans", face = "bold"),
        legend.position = c(0.15,0.7 ),
        legend.title = element_blank(),
        legend.text = element_text(size = 12, family = "sans", face = "bold"),
        legend.background =  element_rect(fill = "transparent", colour = "transparent"),
        legend.key = element_rect(fill = "transparent", colour = "transparent"),
        legend.key.size = unit(0.15, "cm"),
        legend.key.width = unit(0.2, "cm"),
        legend.spacing.y = unit(-0.1, "cm"),
        panel.background = element_rect(fill = "transparent"),
        plot.background = element_rect(fill = "transparent", color = NA)) +
  scale_y_continuous(limits = c(0,610), expand = c(0,1)) +
  scale_x_discrete(breaks = c("FF1Bov", "FF2Bov", "FF3Bov", "FF1Hum", "FF2Hum", "FF3Hum", "FF4Hum", "FF5Hum", "FF6Hum", "FF7Hum"),
                   labels = c("Bov 1", "Bov 2", "Bov 3", "Hum 1", "Hum 2", "Hum 3", "Hum 4", "Hum 5", "Hum 6", "Hum 7")) +
  xlab(expression(bold(Follicular~fluid~sample))) +
  ylab(expression(bold(Number~of~particles~per~mL)))

C
```

Reproduction of Fig. 1C, composition of plastic particles found in cow
follicular fluid samples as well as the water controls.

```
C_Inset <- 
  ggplot(data = data2[-which(data2$Sample == "FF4Hum"),], 
         aes(x = Sample,
             y = particles_ml,
             fill = Microplastic)) +
  geom_bar(stat = "identity", alpha = 0.8) +
  scale_fill_manual(labels = c("ABS",
                               "Cellulose",
                               "Nylon",
                               "Other",
                               "Polyamide",
                               "Polyester",
                               "Polyethylene",
                               "Polypropoylene",
                               "Polystyrene",
                               "Polyurethene",
                               "Polyvinyl chloride",
                               "Rubber",
                               "Silicon(IV) oxide"),
                    values = c("#001219", "#8d96a3", "#005F73", "#0A9396", "#94D2BD",
                               "#E9D8A6","#EE9B00", "#CA6702",  "#9B2226",  "#6d597a", "#355070",  "#fbb13c", "#bf0603")) +
  theme_bw()+
  theme(panel.grid.major = element_blank(),
        panel.grid.minor = element_blank(),
        axis.title.y = element_text(size=14, family = "sans", face = "bold"),
        axis.title.x = element_text(size=14, family = "sans", face = "bold"),
        axis.text.y = element_text(size=12, family = "sans"),
        axis.text.x  = element_text(size=12, family = "sans"),
        plot.title = element_text(hjust = -0.05, size = 10, family = "sans", face = "bold"),
        legend.position = c(0.15,0.7 ),
        legend.title = element_blank(),
        legend.text = element_text(size = 10, family = "sans", face = "bold"),
        legend.background =  element_rect(fill = "transparent", colour = "transparent"),
        legend.key = element_rect(fill = "transparent", colour = "transparent"),
        legend.key.size = unit(0.15, "cm"),
        legend.key.width = unit(0.2, "cm"),
        legend.spacing.y = unit(-0.1, "cm"),
        panel.background = element_rect(fill = "transparent"),
        plot.background = element_rect(fill = "transparent", color = NA)) +
  scale_y_continuous(limits = c(0,130), expand = c(0,1)) +
  scale_x_discrete(breaks = c("FF1Bov", "FF2Bov", "FF3Bov", "FF1Hum", "FF2Hum", "FF3Hum", "FF4Hum", "FF5Hum", "FF6Hum", "FF7Hum"),
                   labels = c("Bov 1", "Bov 2", "Bov 3", "Hum 1", "Hum 2", "Hum 3", "Hum 4", "Hum 5", "Hum 6", "Hum 7")) +
  xlab(expression(bold(Follicular~fluid~sample))) +
  ylab(expression(bold(Number~of~particles~per~mL)))

C_Inset
```

Reproduction of Fig. 1C, composition of plastic particles found in cow
follicular fluid samples as well as the water controls.

```
D <- 
  ggplot(data = data, 
         aes(x = Sample,
             y = Count,
             fill = Classification)) +
  geom_bar(position = "fill", stat = "identity", alpha = 0.8) +
  scale_fill_manual(labels = c("Coating, solvent, or filler",
                               "Fiber",
                               "Non-plastic related",
                               "Pigment",
                               "Plastic polymer",
                               "Plasticizer"),
                    values = c("#00798c", "#d1495b", "#edae49", "#66a182", "#2e4057", "#8d96a3")) +
  theme_bw()+
  theme(panel.grid.major = element_blank(),
        panel.grid.minor = element_blank(),
        axis.title.y = element_text(size=14, family = "sans", face = "bold"),
        axis.title.x = element_text(size=14, family = "sans", face = "bold"),
        axis.text.y = element_text(size=12, family = "sans"),
        axis.text.x  = element_text(size=10, family = "sans"),
        plot.title = element_text(hjust = -0.05, size = 10, family = "sans", face = "bold"),
        legend.position = "top",
        legend.title = element_blank(),
        legend.text = element_text(size = 9, family = "sans", face = "bold"),
        legend.background =  element_rect(fill = "transparent", colour = "transparent"),
        legend.key = element_rect(fill = "transparent", colour = "transparent"),
        legend.key.size = unit(0.15, "cm"),
        legend.key.width = unit(0.2, "cm"),
        legend.spacing.y = unit(-0.1, "cm"),
        panel.background = element_rect(fill = "transparent"),
        plot.background = element_rect(fill = "transparent", color = NA)) +
  scale_y_continuous(expand = c(0,0.005)) +
  scale_x_discrete(breaks = c("FF1bov", "FF2bov", "FF3bov", "FF1hum", "FF2hum", "FF3hum", "FF4hum", "FF5hum", "FF6hum", "FF7hum", "WtCTRL1", "WtCTRL2", "WtCTRL3", "WtCTRL4", "WtCTRL5"),
                   labels = c("Bov 1", "Bov 2", "Bov 3", "Hum 1", "Hum 2", "Hum 3", "Hum 4", "Hum 5", "Hum 6", "Hum 7", "CTRL 1", "CTRL 2", "CTRL 3", "CTRL 4", "CTRL 5")) +
  xlab(expression(bold(Sample))) +
  ylab(expression(bold(Proportion~of~particles~per~mL)))

D
```

Reproduction of Fig. 1D Bar chart depicting the number of plastic
particles found in cow follicular fluid samples.

#### MPs reduce oocyte maturation and induce damage of the zona pellucidae

The first step in this anaylsis was to import the oocyte maturation
data and carry out some basic data carpentry in order to get the data in
the correct format for analysis.

```
# Import the oocyte maturation data
data <- read.csv("Data/Oocyte_data.csv")

# Scale percentages between 0-1 for the binomial model
data$Percentage2 <- data$Percentage/100

# Rename the groups
data[which(data$Treatment == "CT"),"Treatment"] <- "Control"
data[which(data$Treatment == "B0.3"),"Treatment"] <- "MP_0.3"
data[which(data$Treatment == "B1.1"),"Treatment"] <- "MP_1.1"
data[which(data$Treatment == "B0.05"),"Treatment"] <- "MP_0.05"
data[which(data$Treatment == "B0.1"),"Treatment"] <- "MP_0.1"

#Convert to factors
data$Treatment <- as.factor(data$Treatment)
data$Nuclear_stage <- factor(data$Nuclear_stage, ordered = T, levels = c("BZP", "DG", "MT"))
data$replicate <- as.factor(data$Replicate)
```

After importing and preparing the data, the next step was to obtain
some basic summary statistics.

```
# Mean percentage of oocytes by treatment and nuclear stage
means <- aggregate(Percentage2 ~ Nuclear_stage*Treatment, data = data, FUN = "mean")

# SDs by treatment and stage
sds <- aggregate(Percentage2 ~ Nuclear_stage*Treatment, data = data, FUN = "sd")

means$SD <- sds$Percentage2
names(means)[3] <- c("Mean")

kable(means, format="markdown", digits = 3, align = "c", caption = "Table S1.1 Basic summary statistics on the oocyte maturation data. BZP refers to oocytes with broken zone pellucidae, DG to degenerated oocytes, and MT to mature oocytes.")
```

Table S1.1 Basic summary statistics on the oocyte maturation
data. BZP refers to oocytes with broken zone pellucidae, DG to
degenerated oocytes, and MT to mature oocytes.

| Nuclear\_stage | Treatment | Mean | SD |
| --- | --- | --- | --- |
| BZP | Control | 0.044 | 0.052 |
| DG | Control | 0.275 | 0.123 |
| MT | Control | 0.681 | 0.103 |
| BZP | MP\_0.3 | 0.214 | 0.076 |
| DG | MP\_0.3 | 0.366 | 0.149 |
| MT | MP\_0.3 | 0.420 | 0.148 |
| BZP | MP\_1.1 | 0.262 | 0.109 |
| DG | MP\_1.1 | 0.328 | 0.152 |
| MT | MP\_1.1 | 0.410 | 0.176 |

After an initial visual verification, we found that we needed to
assess whether there was any evidence of under/overdispersion in the
oocyte maturation data. This was done by estimating the dispersion
parameter of the data using a generalised linear regression model with a
quasi-binomial distribution.

```
#Test if the data are under/overdispersed
quasi_binom <- glm(Percentage2 ~ Treatment + Nuclear_stage, data = data, family=quasibinomial)
summary(quasi_binom)
```

```
## 
## Call:
## glm(formula = Percentage2 ~ Treatment + Nuclear_stage, family = quasibinomial, 
##     data = data)
## 
## Deviance Residuals: 
##      Min        1Q    Median        3Q       Max  
## -0.61689  -0.21810   0.00218   0.23649   0.59132  
## 
## Coefficients:
##                   Estimate Std. Error t value Pr(>|t|)    
## (Intercept)     -7.627e-01  2.192e-01  -3.479  0.00152 ** 
## TreatmentMP_0.3  2.451e-04  3.071e-01   0.001  0.99937    
## TreatmentMP_1.1  2.619e-12  3.071e-01   0.000  1.00000    
## Nuclear_stage.L  1.116e+00  2.298e-01   4.855 3.26e-05 ***
## Nuclear_stage.Q -2.715e-02  2.166e-01  -0.125  0.90107    
## ---
## Signif. codes:  0 '***' 0.001 '**' 0.01 '*' 0.05 '.' 0.1 ' ' 1
## 
## (Dispersion parameter for quasibinomial family taken to be 0.115423)
## 
##     Null deviance: 6.9459  on 35  degrees of freedom
## Residual deviance: 3.9141  on 31  degrees of freedom
## AIC: NA
## 
## Number of Fisher Scoring iterations: 4
```

These results indicate that the dispersion parameter was 0.115. This
suggested that the data were underdispersed, requiring that we use the
quasi-binomial distribution when modelling the oocyte maturation
data.

##### Mature oocytes

We used hierarchical generalised linear regression models to assess
the effects of MP exposure on the percentage of mature oocytes after a
24 hour maturation period. For these analyses all models had a
quasi-binomial error distribution to account for the fact that the data
were underdispersed percentages (detailed above), which were rescaled to
range between 0 – 1. We also included random intercepts to account for
any differences across replicates.

```
MT.fit <- glmmPQL(Percentage2 ~ Treatment,
                  random = ~ 1|Replicate,
                  family=quasibinomial,
                  data = data[which(data$Nuclear_stage == "MT"),])

summary(MT.fit)
```

```
## Linear mixed-effects model fit by maximum likelihood
##   Data: data[which(data$Nuclear_stage == "MT"), ] 
##   AIC BIC logLik
##    NA  NA     NA
## 
## Random effects:
##  Formula: ~1 | Replicate
##         (Intercept)  Residual
## StdDev:   0.4982103 0.1177631
## 
## Variance function:
##  Structure: fixed weights
##  Formula: ~invwt 
## Fixed effects:  Percentage2 ~ Treatment 
##                      Value Std.Error DF   t-value p-value
## (Intercept)      0.7966665 0.3245090  6  2.454990  0.0495
## TreatmentMP_0.3 -1.1398080 0.2068783  6 -5.509557  0.0015
## TreatmentMP_1.1 -1.1849105 0.2072735  6 -5.716653  0.0012
##  Correlation: 
##                 (Intr) TMP_0.
## TreatmentMP_0.3 -0.338       
## TreatmentMP_1.1 -0.337  0.533
## 
## Standardized Within-Group Residuals:
##        Min         Q1        Med         Q3        Max 
## -1.5253196 -0.5435709 -0.1934040  0.5956740  1.4625857 
## 
## Number of Observations: 12
## Number of Groups: 4
```

```
# Tukey post-hoc test
MT.TUKEY <- glht(MT.fit,
                 mcp(Treatment = "Tukey"),
                 data = data[which(data$Nuclear_stage == "MT"),])
summary(MT.TUKEY)
```

```
## 
##   Simultaneous Tests for General Linear Hypotheses
## 
## Multiple Comparisons of Means: Tukey Contrasts
## 
## 
## Fit: glmmPQL(fixed = Percentage2 ~ Treatment, random = ~1 | Replicate, 
##     family = quasibinomial, data = data[which(data$Nuclear_stage == 
##         "MT"), ])
## 
## Linear Hypotheses:
##                       Estimate Std. Error z value Pr(>|z|)    
## MP_0.3 - Control == 0  -1.1398     0.1792  -6.362   <1e-05 ***
## MP_1.1 - Control == 0  -1.1849     0.1795  -6.601   <1e-05 ***
## MP_1.1 - MP_0.3 == 0   -0.0451     0.1734  -0.260    0.963    
## ---
## Signif. codes:  0 '***' 0.001 '**' 0.01 '*' 0.05 '.' 0.1 ' ' 1
## (Adjusted p values reported -- single-step method)
```

These results indicate that exposure to MPs resulted in significantly
reduced oocyte maturation rates compared to the control. A Tukey
post-hoc comparison found that while exposure to both the 0.3 \(\mu\mathrm{m}\) and 1.1 \(\mu\mathrm{m}\) MP beads resulted in
significantly lower maturation rates than the control, there was no
significant difference between the two sized beads.

##### Degraded cells

We used hierarchical generalised linear regression models to assess
the effects of MP exposure on the percentage of degraded oocytes after a
24 hour maturation period. For these analyses all models had a
quasi-binomial error distribution to account for the fact that the data
were underdispersed percentages, ranging between 0 – 1. We included
random intercepts to account for any differences across replicates.

```
DG.fit <- glmmPQL(Percentage2 ~ Treatment,
                  random = ~ 1|Replicate,
                  family=quasibinomial,
                  data = data[which(data$Nuclear_stage == "DG"),])

summary(DG.fit)
```

```
## Linear mixed-effects model fit by maximum likelihood
##   Data: data[which(data$Nuclear_stage == "DG"), ] 
##   AIC BIC logLik
##    NA  NA     NA
## 
## Random effects:
##  Formula: ~1 | Replicate
##         (Intercept)  Residual
## StdDev:   0.4841533 0.1373825
## 
## Variance function:
##  Structure: fixed weights
##  Formula: ~invwt 
## Fixed effects:  Percentage2 ~ Treatment 
##                      Value Std.Error DF   t-value p-value
## (Intercept)     -1.0163720 0.3338075  6 -3.044785  0.0227
## TreatmentMP_0.3  0.4414409 0.2479213  6  1.780568  0.1253
## TreatmentMP_1.1  0.2655144 0.2507173  6  1.059019  0.3304
##  Correlation: 
##                 (Intr) TMP_0.
## TreatmentMP_0.3 -0.400       
## TreatmentMP_1.1 -0.395  0.531
## 
## Standardized Within-Group Residuals:
##          Min           Q1          Med           Q3          Max 
## -1.310939697 -0.618900428  0.008632261  0.521490568  1.799081925 
## 
## Number of Observations: 12
## Number of Groups: 4
```

```
# Tukey post-hoc test
DG.TUKEY <- glht(DG.fit,
                 mcp(Treatment = "Tukey"),
                 data = data[which(data$Nuclear_stage == "DG"),])
summary(DG.TUKEY)
```

```
## 
##   Simultaneous Tests for General Linear Hypotheses
## 
## Multiple Comparisons of Means: Tukey Contrasts
## 
## 
## Fit: glmmPQL(fixed = Percentage2 ~ Treatment, random = ~1 | Replicate, 
##     family = quasibinomial, data = data[which(data$Nuclear_stage == 
##         "DG"), ])
## 
## Linear Hypotheses:
##                       Estimate Std. Error z value Pr(>|z|)  
## MP_0.3 - Control == 0   0.4414     0.2147   2.056   0.0992 .
## MP_1.1 - Control == 0   0.2655     0.2171   1.223   0.4395  
## MP_1.1 - MP_0.3 == 0   -0.1759     0.2092  -0.841   0.6775  
## ---
## Signif. codes:  0 '***' 0.001 '**' 0.01 '*' 0.05 '.' 0.1 ' ' 1
## (Adjusted p values reported -- single-step method)
```

These results indicate that exposure to MPs tended to result in a
higher percentage of degraded oocyte compared to the control, but the
effects were not statistically significant. A Tukey post-hoc comparison
found no evidence of any pair-wise differences.

##### Broken zona pellucidae

We used generalised linear regression models to assess the effects of
MP exposure on the percentage of oocytes with broken zona pellucidae
after a 24 hour maturation period. For these analyses all models had a
quasi-binomial error distribution to account for the fact that the data
were underdispersed percentages, ranging between 0 – 1. We also included
random intercepts to account for any differences across replicates.

```
BZP.fit <- glmmPQL(Percentage2 ~ Treatment,
                   random = ~ 1|Replicate,
                   family=quasibinomial,
                   data = data[which(data$Nuclear_stage == "BZP"),])

summary(BZP.fit)
```

```
## Linear mixed-effects model fit by maximum likelihood
##   Data: data[which(data$Nuclear_stage == "BZP"), ] 
##   AIC BIC logLik
##    NA  NA     NA
## 
## Random effects:
##  Formula: ~1 | Replicate
##         (Intercept)  Residual
## StdDev:   0.3334861 0.1589298
## 
## Variance function:
##  Structure: fixed weights
##  Formula: ~invwt 
## Fixed effects:  Percentage2 ~ Treatment 
##                     Value Std.Error DF   t-value p-value
## (Intercept)     -3.105634 0.4875155  6 -6.370330  0.0007
## TreatmentMP_0.3  1.782692 0.5006382  6  3.560839  0.0119
## TreatmentMP_1.1  2.053075 0.4941206  6  4.155007  0.0060
##  Correlation: 
##                 (Intr) TMP_0.
## TreatmentMP_0.3 -0.820       
## TreatmentMP_1.1 -0.831  0.808
## 
## Standardized Within-Group Residuals:
##         Min          Q1         Med          Q3         Max 
## -1.39173435 -0.81840743 -0.04819769  0.66010424  1.29451024 
## 
## Number of Observations: 12
## Number of Groups: 4
```

```
# Tukey post-hoc test
BZP.TUKEY <- glht(BZP.fit,
                  mcp(Treatment = "Tukey"),
                  data = data[which(data$Nuclear_stage == "BZP"),])
summary(BZP.TUKEY)
```

```
## 
##   Simultaneous Tests for General Linear Hypotheses
## 
## Multiple Comparisons of Means: Tukey Contrasts
## 
## 
## Fit: glmmPQL(fixed = Percentage2 ~ Treatment, random = ~1 | Replicate, 
##     family = quasibinomial, data = data[which(data$Nuclear_stage == 
##         "BZP"), ])
## 
## Linear Hypotheses:
##                       Estimate Std. Error z value Pr(>|z|)    
## MP_0.3 - Control == 0   1.7827     0.4336   4.112  0.00011 ***
## MP_1.1 - Control == 0   2.0531     0.4279   4.798  < 1e-05 ***
## MP_1.1 - MP_0.3 == 0    0.2704     0.2668   1.014  0.56061    
## ---
## Signif. codes:  0 '***' 0.001 '**' 0.01 '*' 0.05 '.' 0.1 ' ' 1
## (Adjusted p values reported -- single-step method)
```

These results indicate that exposure to MPs resulted in a
significantly higher percentage of oocytes with broken zona pellucidae
after a 24 hour maturation period compared to the control. A Tukey
post-hoc comparison found that while exposure to both the 0.3 \(\mu\mathrm{m}\) and 1.1 \(\mu\mathrm{m}\) MP beads resulted in
significantly more broken zona pellucidae than the control, there was no
significant difference in the number of broken zona pellucidae between
the two size treatments. A figure of the results across all three
functional traits is shown below.

```
A <- 
  ggplot(data = data, 
         aes(x = Nuclear_stage,
             y = Percentage,
             fill = Treatment)) +
  geom_point(aes(col = Treatment),
             position = position_jitterdodge(jitter.width = 0.05),
             size = 0.3) +
  geom_boxplot(size = 0.2,
               alpha = 0.5,
               outlier.size = 0,
               outlier.shape = NA) +
  scale_fill_manual(labels = c("Control",
                               "0.3 \U03BCm",
                               "1.1 \U03BCm"),
                    values = c("black", '#dfb433', '#3c7a47'),
                    guide = "none") +
  scale_colour_manual(labels = c("Control",
                                 "0.3 \U03BCm",
                                 "1.1 \U03BCm"),
                      values = c("black", '#dfb433', '#3c7a47')) +
  theme_bw()+
  theme(panel.grid.major = element_blank(),
        panel.grid.minor = element_blank(),
        axis.title.y = element_text(size=10, family = "sans", face = "bold"),
        axis.title.x = element_blank(),
        axis.text.y = element_text(size=8, family = "sans", face = "bold"),
        axis.text.x  = element_text(size=8, family = "sans"),
        plot.title = element_text(hjust = -0.05, size = 10, family = "sans", face = "bold"),
        legend.position = c(0.1,0.8 ),
        legend.title = element_blank(),
        legend.text = element_text(size = 8, family = "sans", face = "bold"),
        legend.background =  element_rect(fill = "transparent", colour = "transparent"),
        legend.key = element_rect(fill = "transparent", colour = "transparent"),
        legend.key.size = unit(0.15, "cm"),
        legend.key.width = unit(0.2, "cm"),
        legend.spacing.y = unit(-0.1, "cm"),
        panel.background = element_rect(fill = "transparent"),
        plot.background = element_rect(fill = "transparent", color = NA)) +
  scale_y_continuous(limits = c(0,100), expand = c(0,1)) +
  scale_x_discrete(breaks = c("BZP", "DG", "MT"), labels = c("Broken zona pellucidae", "Degraded", "Mature")) +
  ylab(expression(bold(Percentage~of~oocytes~"(%)")))  +
  guides(colour = guide_legend(override.aes = list(size=1))) 

A
```

Reproduction of Fig. 2A Boxplots depicting oocyte maturation over time
as a function of microplastics treatment.

#### MPs alter the oocyte proteome

##### Data normalisation

The first step was to transform the raw data so that they would be on
the same scale. To render the data comparable across species and
developmental stages, data were scaled using Probabilistic Quotient
Normalisation (PQN), which calibrates individual gene expression
profiles against the median profile. Notably, analyses on PQN
transformed data have been shown to have low false-positive rates, and
can accurately recover groups of interest without introducing
artefactual differences.

```
#Import the raw data
data <- read.csv("Data/Proteomics_oocyte_MP_raw.csv")

# Some data carpentry
# Only keep a single one of the multiple IDs
data$Protein.IDs <- sub(";.*", "", data$Protein.IDs)
Proteins <- data$Protein.IDs


#Identify the gene names
GN <- sub(".*GN=", "", data$Fasta.headers)
GN <-  sub(" .*", "", GN)
GN[grepl( 'tr|', GN, fixed = TRUE)] <- NA
data$Gene_Name <- GN

#Store the proteins expressed within the different groups
control <- rowMeans(data[,c("CT_1","CT_2","CT_3","CT_4","CT_5")])
B0.3 <- rowMeans(data[,c("B0.3_1","B0.3_2","B0.3_3","B0.3_4","B0.3_5")])
B1.1 <- rowMeans(data[,c("B1.1_1","B1.1_2","B1.1_3","B1.1_4","B1.1_5")])

#Transpose the data and assign the correct column names
data <- data.frame(t(data[,c(7:21)]))
colnames(data) <- Proteins

#Store the untransformed data in memory
raw_data <- data


# Create vectors with treatment IDs
Treatment <- c("1.1", "1.1", "1.1", "control", "0.3", "0.3", "0.3", "1.1", "control", "control", "0.3", "0.3", "control", "control", "1.1")
Treatment <- as.factor(Treatment)

Treatment_2 <- c("MP", "MP", "MP", "control", "MP", "MP", "MP", "MP", "control", "control", "MP", "MP", "control", "control", "MP")
Treatment_2 <- as.factor(Treatment_2)

###############################################################
# PQN Normalisation of the data
###############################################################

#Calculate the median of each gene's expression to generate a reference sample
ref <- as.vector(apply(data, 2, median))

for(i in 1:nrow(data)){
  QUOTIENTS <- data[i,]/ref
  m_j <- median(t(QUOTIENTS), na.rm = T)
  data[i,] <- data[i,]/m_j
  data[i,][data[i,] == min(data[i,])] <- 0
}
```

##### Random Forest Classification

After the data were transformed, a random forest (RF) model was used
to classify protein expression profiles according to whether or not the
oocytes were exposed to microplastics, with scaled gene expression
values as the prediction variables. This allowed us to determine how
well information contained within the proteomics data could be used to
predict MP exposure. These analyses were conducted using the R package
randomForest. We chose RF modeling as it does not require any parameter
reduction prior to analysis. Identification of genes important for
classifying groups of interest in each RF model was carried out using RF
variable importance values.

```
#Create a dataset that has information on MP exposure vs. controls
Treatment_Data <- data
Treatment_Data$Treatment <- Treatment


#Run the random forest model identifying MP exposure treatments
exposure.mod <- randomForest(y = Treatment_Data$Treatment,
                             x = Treatment_Data[, colnames(Treatment_Data) != "Treatment"],
                             mtry =  5,
                             ntree= 20000,
                             importance=TRUE,
                             proximity = TRUE,
                             keep.forest=TRUE,
                             replace = TRUE)

exposure.mod
```

```
## 
## Call:
##  randomForest(x = Treatment_Data[, colnames(Treatment_Data) !=      "Treatment"], y = Treatment_Data$Treatment, ntree = 20000,      mtry = 5, replace = TRUE, importance = TRUE, proximity = TRUE,      keep.forest = TRUE) 
##                Type of random forest: classification
##                      Number of trees: 20000
## No. of variables tried at each split: 5
## 
##         OOB estimate of  error rate: 86.67%
## Confusion matrix:
##         0.3 1.1 control class.error
## 0.3       0   1       4         1.0
## 1.1       2   1       2         0.8
## control   3   1       1         0.8
```

```
confusionMatrix(exposure.mod$predicted, Treatment_Data$Treatment)
```

```
## Confusion Matrix and Statistics
## 
##           Reference
## Prediction 0.3 1.1 control
##    0.3       0   2       3
##    1.1       1   1       1
##    control   4   2       1
## 
## Overall Statistics
##                                           
##                Accuracy : 0.1333          
##                  95% CI : (0.0166, 0.4046)
##     No Information Rate : 0.3333          
##     P-Value [Acc > NIR] : 0.9806          
##                                           
##                   Kappa : -0.3            
##                                           
##  Mcnemar's Test P-Value : 0.8472          
## 
## Statistics by Class:
## 
##                      Class: 0.3 Class: 1.1 Class: control
## Sensitivity              0.0000    0.20000        0.20000
## Specificity              0.5000    0.80000        0.40000
## Pos Pred Value           0.0000    0.33333        0.14286
## Neg Pred Value           0.5000    0.66667        0.50000
## Prevalence               0.3333    0.33333        0.33333
## Detection Rate           0.0000    0.06667        0.06667
## Detection Prevalence     0.3333    0.20000        0.46667
## Balanced Accuracy        0.2500    0.50000        0.30000
```

```
varImpPlot(exposure.mod, type=1, cex = 0.5, main = "MPs and the oocyte proteome")
```

These results indicate that the RF model was able to classify samples
with versus without MP exposure with an accuracy of ca. 20%, which did
not represent a significant improvement over the null, no-information
rate of classification.

```
res.pca <- PCA(exposure.mod$proximity, graph = FALSE) # Conduct a PCA on the proximity matrix
PC1 <- res.pca$ind$coord[,1] #Store individual coordinates of PC1 as a vector
PC2 <- res.pca$ind$coord[,2] #Store individual coordinates of PC2 as a vector
PCs.ID <- data.frame(cbind(PC1,PC2)) #Bind the coordinates together as a dataframe
PCs.ID$Treatment <- Treatment_Data$Treatment #Add in treatment to the df

#Define axis labels based on % of data explained across each dimension of the PCA
DIM_1 <- paste("Dimension 1 (", round(res.pca$eig[1,2], 1), "%)")
DIM_2 <- paste("Dimension 2 (", round(res.pca$eig[2,2], 1), "%)")

#Draw ellipses around the clusters and highlighting the elipsoids rather than the data points
centroids <- aggregate(cbind(PC1,PC2)~Treatment,PCs.ID,mean)
conf.rgn  <- do.call(rbind,lapply(unique(PCs.ID$Treatment),function(t)
  data.frame(Treatment=as.character(t),
             ellipse(cov(PCs.ID[PCs.ID$Treatment==t,1:2]),
                     centre=as.matrix(centroids[t,2:3]),
                     level=0.95),
             stringsAsFactors=FALSE)))

#Then make the figure
PCA_FIG <- 
  ggplot(PCs.ID, aes(x=PC1, y=PC2, color = Treatment)) +
  geom_hline(aes(yintercept=0), linetype="dashed", lwd = 0.1) +
  geom_vline(aes(xintercept=0), linetype="dashed", lwd = 0.1) +
  geom_path(data=conf.rgn, alpha=0.2, size = 0) +
  geom_polygon(data=conf.rgn,
               aes(fill = Treatment),
               alpha=0.1,
               size = 0.1,
               show.legend = FALSE) + 
  geom_point(size=0.4) +
  theme_bw() +
  ylab(DIM_2) +
  xlab(DIM_1) + 
  geom_point(size=0.4, aes(color = Treatment)) +
  scale_fill_manual(labels = c("0.3 \U03BCm",
                               "1.1 \U03BCm",
                               "Control"),
                    values = c('#dfb433', '#3c7a47',"black"),
                    guide = "none") +
  scale_colour_manual(labels = c("0.3 \U03BCm",
                                 "1.1 \U03BCm",
                                 "Control"),
                      values = c('#dfb433', '#3c7a47',"black")) +
  ylab(DIM_2) +
  xlab(DIM_1) + 
  theme_bw()+
  theme(panel.grid.major = element_blank(),
        panel.grid.minor = element_blank(),
        axis.title.y = element_text(size=10, family = "sans", face = "bold"),
        axis.title.x = element_text(size=10, family = "sans", face = "bold"),
        axis.text.y = element_text(size=8, family = "sans", face = "bold"),
        axis.text.x  = element_text(size=8, family = "sans"),
        plot.title = element_text(hjust = -0.05, size = 10, family = "sans", face = "bold"),
        legend.position = c(0.1,0.92),
        legend.title = element_blank(),
        legend.text = element_text(size = 6, family = "sans", face = "bold"),
        legend.background =  element_rect(fill = "transparent", colour = "transparent"),
        legend.key = element_rect(fill = "transparent", colour = "transparent"),
        legend.key.size = unit(0.15, "cm"),
        legend.key.width = unit(0.2, "cm"),
        legend.spacing.y = unit(-0.1, "cm"),
        panel.background = element_rect(fill = "transparent"),
        plot.background = element_rect(fill = "transparent", color = NA))

PCA_FIG
```

Reproduction of Fig. 2B PCA on the proximity matrix of the RF
classification model.

```
ggsave(PCA_FIG,
       file="Figures/Treatment_PCA.png",
       width = 3.23,
       height=3,
       units = "in",
       dpi = 600)
```

Density plots for the three proteins of primary importance for
predicting which treatment group oocytes were from are shown below. A
Venn diagram depicting the number of proteins common to each treatment
group is also shown. Note: The R code for producing these figures is not
shown.

```
##  [1] "A6QLV3"     "A4FUA8"     "Q2NKZ1"     "A0A3Q1MKJ7" "F1MBE7"    
##  [6] "A0A3Q1LUG9" "F1MPB2"     "Q32LE5"     "Q56JV9"     "Q07130"    
## [11] "Q3SZ20"     "P68301"     "Q3T133"     "G3N3D4"     "A1A4J1"    
## [16] "A0A3Q1LV36" "Q2TBQ5"     "A3KN04"     "Q862S8"     "Q08E34"
```

Figure S1.1 Density plots of the variables of primary importance for
predicting MP exposure.

Figure S1.2 Venn diagram of the proteins expressed within the different
groups.

```
## quartz_off_screen 
##                 2
```

##### Differentially Expressed Proteins (DEPs)

Differentially expressed proteins (DEPs) of normalized data were
identified using the R package `msEmpiRe`.

```
#Convert to a numeric matrix and log transform
DATA <- as.matrix(t(data))
DATA <- log(DATA + 1)
DATA <- DATA[,-4]

#Create the design matrix
design <- cbind(Exposure = as.numeric(Treatment_2)[-4],
                Treatment = as.numeric(Treatment)[-4])

#Run the analysis and compile the results
test <- lmFit(DATA, design = model.matrix(~ 1 + design[,1]))
test2 <- eBayes(test)
test3 <- topTable(test2, number = length(test2$coefficients), sort.by = "none")

write.csv(test3, file = "Results/DEPs.csv")
```

```
data <- read.csv("Results/B0.3_vs_CT_MsEmpire.csv")

#Which proteins are differentially expressed
labs <- data[which(data$p.adj < 0.05),"gene.names"]
labs[10] <- "H2AC16"
coords <- data[which(data$p.adj < 0.05),c("log2FC", "p.adj")]
coords$p.adj <- -log(coords$p.adj)
labs
```

```
##  [1] "MVP"    "PADI6"  "TXN"    "GPX1"   "DBI"    "AK2"    "MDH2"   "SNRPD2"
##  [9] "HSPE1"  "H2AC16" "TPI1"   "NLRP5"  "ENO1"
```

```
Volcano_Plot <- 
  ggplot(data, aes(x=log2FC, y=-log(p.adj), color = ifelse(p.adj < 0.05,"red", "#046C9A"))) +
  geom_point(size=0.4) +
  scale_color_manual(labels=c("Control", "MP exposure"),
                     values = c("#046C9A", "red"),
                     guide = "none") +
  theme_bw()+
  theme(panel.grid.major = element_blank(),
        panel.grid.minor = element_blank(),
        axis.title.y = element_text(size=14, family = "sans", face = "bold"),
        axis.title.x = element_text(size=14, family = "sans", face = "bold"),
        axis.text.y = element_text(size=12, family = "sans", face = "bold"),
        axis.text.x  = element_text(size=12, family = "sans"),
        plot.title = element_text(hjust = -0.05, size = 10, family = "sans", face = "bold"),
        legend.position = c(0.15,0.1),
        legend.title = element_blank(),
        legend.text = element_text(size = 6, family = "sans", face = "bold"),
        legend.background =  element_rect(fill = "transparent", colour = "transparent"),
        legend.key = element_rect(fill = "transparent", colour = "transparent"),
        legend.key.size = unit(0.15, "cm"),
        legend.key.width = unit(0.2, "cm"),
        legend.spacing.y = unit(-0.1, "cm"),
        panel.background = element_rect(fill = "transparent"),
        plot.background = element_rect(fill = "transparent", color = NA)) +
  scale_y_continuous(expand = c(0,0.2)) +
  xlab(expression(bold(log[2]~fold~change))) +
  ylab(expression(bold(-log~adjusted~p~value))) +
  geom_text_repel(data = coords, aes(x = log2FC, 
                                     y = p.adj, 
                                     label = labs), col = "red", size = 2)

Volcano_Plot
```

Reproduction of Fig. 2C Volcano plot of the proteomics analysis
comparing oocytes exposed to 0.3 μm MPs vs the controls.

```
ggsave(Volcano_Plot,
       file="Figures/Volcano_03.png",
       width = 5.75,
       height=4,
       units = "in",
       dpi = 600)
```

```
data <- read.csv("Results/B1.1_vs_CT_MsEmpire.csv")

#Which proteins are differentially expressed
labs <- data[which(data$p.adj < 0.05),"gene.names"]
coords <- data[which(data$p.adj < 0.05),c("log2FC", "p.adj")]
coords$p.adj <- -log(coords$p.adj)
labs
```

```
##  [1] "DDX5"     "DSP"      "SFPQ"     "BCLAF1"   "ATP5IF1"  "TBCA"    
##  [7] "DSG1"     "TPM1"     "MVP"      "HNRNPM"   "PADI6"    "NLRP5"   
## [13] "HSP90AA1"
```

```
Volcano_Plot <- 
  ggplot(data, aes(x=log2FC, y=-log(p.adj), color = ifelse(p.adj < 0.05,"red", "#046C9A"))) +
  geom_point(size=0.4) +
  scale_color_manual(labels=c("Control", "MP exposure"),
                     values = c("#046C9A", "red"),
                     guide = "none") +
  theme_bw()+
  theme(panel.grid.major = element_blank(),
        panel.grid.minor = element_blank(),
        axis.title.y = element_text(size=14, family = "sans", face = "bold"),
        axis.title.x = element_text(size=14, family = "sans", face = "bold"),
        axis.text.y = element_text(size=12, family = "sans", face = "bold"),
        axis.text.x  = element_text(size=12, family = "sans"),
        plot.title = element_text(hjust = -0.05, size = 10, family = "sans", face = "bold"),
        legend.position = c(0.15,0.1),
        legend.title = element_blank(),
        legend.text = element_text(size = 6, family = "sans", face = "bold"),
        legend.background =  element_rect(fill = "transparent", colour = "transparent"),
        legend.key = element_rect(fill = "transparent", colour = "transparent"),
        legend.key.size = unit(0.15, "cm"),
        legend.key.width = unit(0.2, "cm"),
        legend.spacing.y = unit(-0.1, "cm"),
        panel.background = element_rect(fill = "transparent"),
        plot.background = element_rect(fill = "transparent", color = NA)) +
  scale_y_continuous(expand = c(0,0.2)) +
  xlab(expression(bold(log[2]~fold~change))) +
  ylab(expression(bold(-log~adjusted~p~value))) +
  geom_text_repel(data = coords, aes(x = log2FC, 
                                     y = p.adj, 
                                     label = labs), col = "red", size = 2)

Volcano_Plot
```

Reproduction of Fig. 2D Volcano plot of the proteomics analysis
comparing oocytes exposed to 1.1 μm MPs vs the controls.

```
ggsave(Volcano_Plot,
       file="Figures/Volcano_11.png",
       width = 5.75,
       height=4,
       units = "in",
       dpi = 600)
```

#### Size distribution in blanks

```
data <- read.csv("Data/Sizes_PP.csv")
data$Sample <- as.factor(data$Sample)


png(filename="Figures/size_L_hum_hist.png",
    width = 6.86, height = 4, units = "in",
    res = 600)

ggplot(data, aes(x=Length, fill=Sample)) +
  geom_histogram(binwidth=.5, alpha=.5, position="identity") + 
  theme_bw() +
  scale_fill_viridis_d(breaks = c("hFF1", "hFF2", "hFF3", "hFF4", "hFF5", "hFF6", "hFF7"),
                       labels = c("Hum 1", "Hum 2", "Hum 3", "Hum 4", "Hum 5", "Hum 6", "Hum 7")) +
  theme(panel.grid.major = element_blank(),
        panel.grid.minor = element_blank(),
        axis.title.y = element_text(size=14, family = "sans", face = "bold"),
        axis.title.x = element_text(size=14, family = "sans", face = "bold"),
        axis.text.y = element_text(size=10, family = "sans", face = "bold"),
        axis.text.x  = element_text(size=12, family = "sans"),
        plot.title = element_text(hjust = -0.05, size = 10, family = "sans", face = "bold"),
        legend.position = "top",
        legend.title = element_blank(),
        legend.text = element_text(size = 6, family = "sans", face = "bold"),
        legend.background =  element_rect(fill = "transparent", colour = "transparent"),
        legend.key = element_rect(fill = "transparent", colour = "transparent"),
        
        panel.background = element_rect(fill = "transparent"),
        plot.background = element_rect(fill = "transparent", color = NA)) +
  scale_x_continuous(limits = c(0,140), expand = c(0,0)) +
  scale_y_continuous(limits = c(0,70), expand = c(0,0)) +
xlab("Length (\U03BCm)")

dev.off()
```

```
## quartz_off_screen 
##                 2
```

```
png(filename="Figures/size_W_hum_Hist.png",
    width = 6.86, height = 4, units = "in",
    res = 600)

ggplot(data, aes(x=Width, fill=Sample)) +
  geom_histogram(binwidth=.5, alpha=.5, position="identity") + 
  theme_bw() +
  scale_fill_viridis_d(breaks = c("hFF1", "hFF2", "hFF3", "hFF4", "hFF5", "hFF6", "hFF7"),
                       labels = c("Hum 1", "Hum 2", "Hum 3", "Hum 4", "Hum 5", "Hum 6", "Hum 7")) +
  theme(panel.grid.major = element_blank(),
        panel.grid.minor = element_blank(),
        axis.title.y = element_text(size=14, family = "sans", face = "bold"),
        axis.title.x = element_text(size=14, family = "sans", face = "bold"),
        axis.text.y = element_text(size=10, family = "sans", face = "bold"),
        axis.text.x  = element_text(size=12, family = "sans"),
        plot.title = element_text(hjust = -0.05, size = 10, family = "sans", face = "bold"),
        legend.position = "top",
        legend.title = element_blank(),
        legend.text = element_text(size = 6, family = "sans", face = "bold"),
        legend.background =  element_rect(fill = "transparent", colour = "transparent"),
        legend.key = element_rect(fill = "transparent", colour = "transparent"),
        
        panel.background = element_rect(fill = "transparent"),
        plot.background = element_rect(fill = "transparent", color = NA)) +
  scale_x_continuous(limits = c(0,50), expand = c(0,0)) +
  scale_y_continuous(limits = c(0,150), expand = c(0,0)) +
xlab("Width (\U03BCm)")

dev.off()
```

```
## quartz_off_screen 
##                 2
```

```
data2 <- read.csv("Data/Sizes_PP_bov.csv")
data2$Sample <- as.factor(data2$Sample)


png(filename="Figures/size_L_bov_Hist.png",
    width = 6.86, height = 4, units = "in",
    res = 600)

ggplot(data2, aes(x=Length, fill=Sample)) +
  geom_histogram(binwidth=.5, alpha=.5, position="identity") + 
  theme_bw() +
  scale_fill_viridis_d(breaks = c("bFF1", "bFF2", "bFF3"),
                       labels = c("Bov 1", "Bov 2", "Bov 3")) +
  theme(panel.grid.major = element_blank(),
        panel.grid.minor = element_blank(),
        axis.title.y = element_text(size=14, family = "sans", face = "bold"),
        axis.title.x = element_text(size=14, family = "sans", face = "bold"),
        axis.text.y = element_text(size=10, family = "sans", face = "bold"),
        axis.text.x  = element_text(size=12, family = "sans"),
        plot.title = element_text(hjust = -0.05, size = 10, family = "sans", face = "bold"),
        legend.position = "top",
        legend.title = element_blank(),
        legend.text = element_text(size = 6, family = "sans", face = "bold"),
        legend.background =  element_rect(fill = "transparent", colour = "transparent"),
        legend.key = element_rect(fill = "transparent", colour = "transparent"),
        
        panel.background = element_rect(fill = "transparent"),
        plot.background = element_rect(fill = "transparent", color = NA)) +
  scale_x_continuous(limits = c(0,52), expand = c(0,0)) +
  scale_y_continuous(limits = c(0,10), expand = c(0,0)) +
xlab("Length (\U03BCm)")

dev.off()
```

```
## quartz_off_screen 
##                 2
```

```
png(filename="Figures/size_W_bov_Hist.png",
    width = 6.86, height = 4, units = "in",
    res = 600)

ggplot(data2, aes(x=Width, fill=Sample)) +
  geom_histogram(binwidth=.5, alpha=.5, position="identity") + 
  theme_bw() +
  scale_fill_viridis_d(breaks = c("bFF1", "bFF2", "bFF3"),
                       labels = c("Bov 1", "Bov 2", "Bov 3")) +
  theme(panel.grid.major = element_blank(),
        panel.grid.minor = element_blank(),
        axis.title.y = element_text(size=14, family = "sans", face = "bold"),
        axis.title.x = element_text(size=14, family = "sans", face = "bold"),
        axis.text.y = element_text(size=10, family = "sans", face = "bold"),
        axis.text.x  = element_text(size=12, family = "sans"),
        plot.title = element_text(hjust = -0.05, size = 10, family = "sans", face = "bold"),
        legend.position = "top",
        legend.title = element_blank(),
        legend.text = element_text(size = 6, family = "sans", face = "bold"),
        legend.background =  element_rect(fill = "transparent", colour = "transparent"),
        legend.key = element_rect(fill = "transparent", colour = "transparent"),
        
        panel.background = element_rect(fill = "transparent"),
        plot.background = element_rect(fill = "transparent", color = NA)) +
  scale_x_continuous(limits = c(0,32), expand = c(0,0)) +
  scale_y_continuous(limits = c(0,18), expand = c(0,0)) +
xlab("Width (\U03BCm)")

dev.off()
```

```
## quartz_off_screen 
##                 2
```

### Technical information

Below the information on the operating system and package versions
for all of the packages used in carrying out these analyses are
detailed.

```
sessionInfo()
```

```
## R version 4.2.1 (2022-06-23)
## Platform: x86_64-apple-darwin17.0 (64-bit)
## Running under: macOS Big Sur ... 10.16
## 
## Matrix products: default
## BLAS:   /Library/Frameworks/R.framework/Versions/4.2/Resources/lib/libRblas.0.dylib
## LAPACK: /Library/Frameworks/R.framework/Versions/4.2/Resources/lib/libRlapack.dylib
## 
## locale:
## [1] en_US.UTF-8/en_US.UTF-8/en_US.UTF-8/C/en_US.UTF-8/en_US.UTF-8
## 
## attached base packages:
## [1] grid      stats     graphics  grDevices utils     datasets  methods  
## [8] base     
## 
## other attached packages:
##  [1] limma_3.52.4         Biobase_2.56.0       BiocGenerics_0.42.0 
##  [4] msEmpiRe_0.1.0       forcats_0.5.2        stringr_1.5.0       
##  [7] dplyr_1.0.10         purrr_1.0.0          readr_2.1.3         
## [10] tidyr_1.2.1          tibble_3.1.8         tidyverse_1.3.2     
## [13] gridExtra_2.3        ggridges_0.5.4       caret_6.0-93        
## [16] lattice_0.20-45      randomForest_4.7-1.1 ellipse_0.4.3       
## [19] FactoMineR_2.7       VennDiagram_1.7.3    futile.logger_1.4.3 
## [22] ggrepel_0.9.2        knitr_1.41           ggplot2_3.4.0       
## [25] gtsummary_1.6.3      MuMIn_1.47.1         multcomp_1.4-20     
## [28] TH.data_1.1-1        survival_3.4-0       mvtnorm_1.1-3       
## [31] lme4_1.1-31          Matrix_1.5-3         MASS_7.3-58.1       
## 
## loaded via a namespace (and not attached):
##   [1] readxl_1.4.1         backports_1.4.1      systemfonts_1.0.4   
##   [4] plyr_1.8.8           splines_4.2.1        listenv_0.9.0       
##   [7] digest_0.6.31        foreach_1.5.2        htmltools_0.5.4     
##  [10] fansi_1.0.3          magrittr_2.0.3       googlesheets4_1.0.1 
##  [13] cluster_2.1.4        tzdb_0.3.0           recipes_1.0.3       
##  [16] globals_0.16.2       modelr_0.1.10        gower_1.0.1         
##  [19] sandwich_3.0-2       hardhat_1.2.0        timechange_0.1.1    
##  [22] colorspace_2.0-3     rvest_1.0.3          textshaping_0.3.6   
##  [25] haven_2.5.1          xfun_0.36            crayon_1.5.2        
##  [28] jsonlite_1.8.4       zoo_1.8-11           iterators_1.0.14    
##  [31] glue_1.6.2           gtable_0.3.1         gargle_1.2.1        
##  [34] ipred_0.9-13         emmeans_1.8.3        future.apply_1.10.0 
##  [37] scales_1.2.1         futile.options_1.0.1 DBI_1.1.3           
##  [40] Rcpp_1.0.9           viridisLite_0.4.1    xtable_1.8-4        
##  [43] proxy_0.4-27         flashClust_1.01-2    stats4_4.2.1        
##  [46] lava_1.7.0           prodlim_2019.11.13   DT_0.26             
##  [49] htmlwidgets_1.6.0    httr_1.4.4           ellipsis_0.3.2      
##  [52] farver_2.1.1         pkgconfig_2.0.3      nnet_7.3-18         
##  [55] multcompView_0.1-8   sass_0.4.4           dbplyr_2.2.1        
##  [58] utf8_1.2.2           labeling_0.4.2       tidyselect_1.2.0    
##  [61] rlang_1.0.6          reshape2_1.4.4       munsell_0.5.0       
##  [64] cellranger_1.1.0     tools_4.2.1          cachem_1.0.6        
##  [67] cli_3.5.0            generics_0.1.3       broom_1.0.2         
##  [70] evaluate_0.19        fastmap_1.1.0        ragg_1.2.4          
##  [73] yaml_2.3.6           ModelMetrics_1.2.2.2 fs_1.5.2            
##  [76] future_1.30.0        nlme_3.1-161         formatR_1.13        
##  [79] leaps_3.1            xml2_1.3.3           compiler_4.2.1      
##  [82] rstudioapi_0.14      e1071_1.7-12         gt_0.8.0            
##  [85] reprex_2.0.2         broom.helpers_1.10.0 bslib_0.4.2         
##  [88] stringi_1.7.8        highr_0.10           nloptr_2.0.3        
##  [91] vctrs_0.5.1          pillar_1.8.1         lifecycle_1.0.3     
##  [94] jquerylib_0.1.4      estimability_1.4.1   data.table_1.14.6   
##  [97] R6_2.5.1             parallelly_1.33.0    codetools_0.2-18    
## [100] lambda.r_1.2.4       boot_1.3-28.1        assertthat_0.2.1    
## [103] withr_2.5.0          parallel_4.2.1       hms_1.1.2           
## [106] rpart_4.1.19         timeDate_4021.107    coda_0.19-4         
## [109] class_7.3-20         minqa_1.2.5          rmarkdown_2.19      
## [112] googledrive_2.0.0    pROC_1.18.0          scatterplot3d_0.3-42
## [115] lubridate_1.9.0
```
