## Supplementary material for "Microplastics are present in women’s and cows’ follicular fluid and polystyrene microplastics compromise bovine oocyte function *in vitro*": Suplementary figures 1 and 2 and tables 1 and 2


**This file includes:**

Supplementary Figures 1 and 2
Supplementary Tables 1 and 2

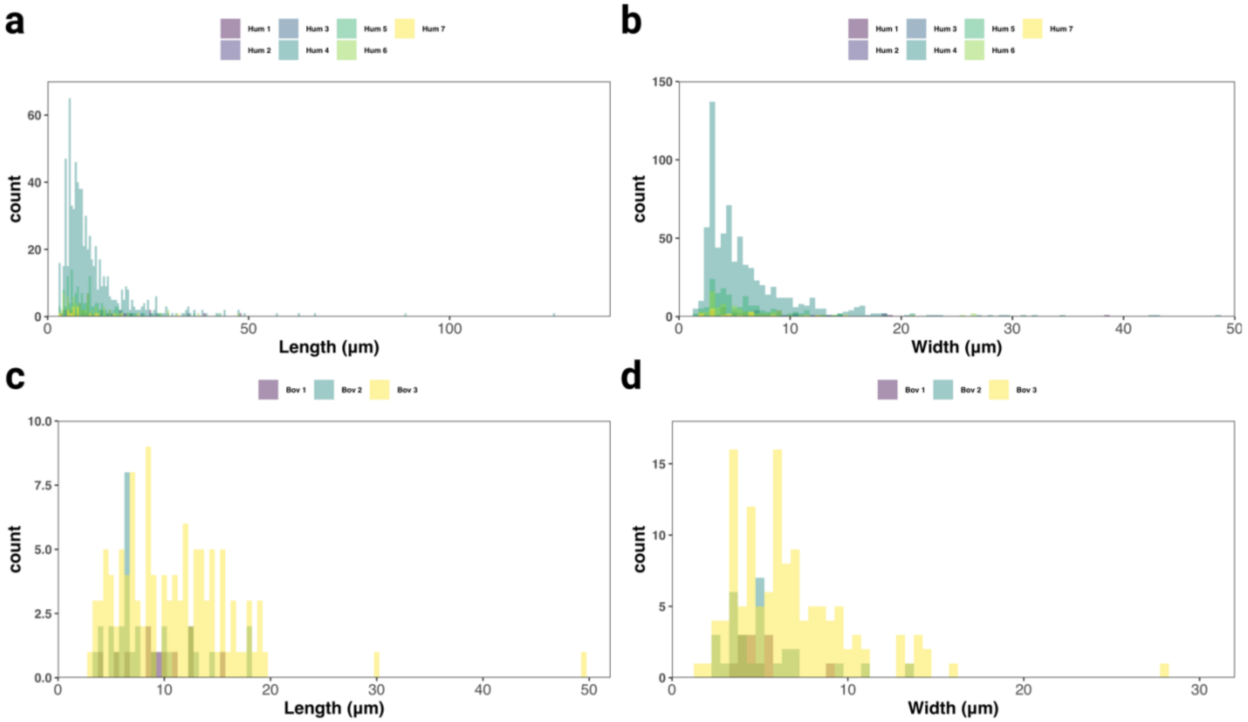


**Supplementary Figure 1. Size distribution of MPs.** Length and width distribution of MPs in human (**a** and **b**) and Bovine (**c** and **d**) follicular fluid samples.

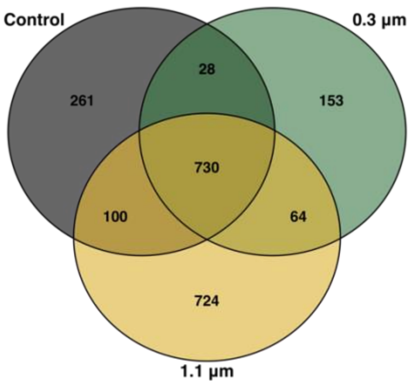


**Supplementary Figure 2. Venn diagram depicting the proteins count distribution of oocytes incubated or not with MPs.** Unique and shared proteins counts are shown for oocytes incubated with 0.3 µm polystyrene beads (0.3 µm), oocytes incubated with 1.1 µm polystyrene beads (1.1 µm) or oocytes that were not exposed to MPs (Control).

**Supplementary Table 1.** Total number of polymers (particles $mL^{-1}$) identified in bovine and human follicular fluid (FF) after adjusting to water controls.

| | Human | | | | | | | Bovine | | |
|---|---|---|---|---|---|---|---|---|---|---|
| | **FF1** | **FF2** | **FF3** | **FF4** | **FF5** | **FF6** | **FF7** | **FF1** | **FF2** | **FF3** |
| ABS | 0.8 | 0.0 | 0.0 | 52.5 | 0 | 0.8 | 0.8 | 0.0 | 0.0 | 1.3 |
| CELLULOSE | 0.0 | 0.0 | 0.0 | 0.0 | 2.5 | 4.2 | 1.7 | 1.3 | 0.4 | 10.4 |
| NYLON | 0.0 | 0.0 | 0.0 | 10.0 | 5.8 | 4.2 | 0 | 0.0 | 0.0 | 1.7 |
| POLYPROPYLENE | 9.2 | 0.0 | 0.0 | 12.5 | 3.3 | 1.7 | 0.8 | 0.0 | 0.0 | 9.6 |
| POLYAMIDE | 0.0 | 0.0 | 0.0 | 1.7 | 5.8 | 9.2 | 2.5 | 0.0 | 0.0 | 0.8 |
| POLYESTER | 0.0 | 0.0 | 0.0 | 10.0 | 2.5 | 5.8 | 0 | 0.0 | 0.0 | 1.7 |
| POLYETHYLENE | 0.0 | 0.8 | 0.0 | 5.8 | 56.7 | 8.3 | 0.8 | 0.0 | 0.0 | 17.1 |
| POLYSTYRENE | 4.2 | 0.0 | 0.0 | 40.0 | 0.8 | 0 | 0 | 0.0 | 1.3 | 9.6 |
| POLYURETHANE | 0.0 | 0.0 | 0.8 | 0.0 | 0 | 0 | 0 | 0.0 | 0.0 | 0.0 |
| POLYVINYL CHLORIDE | 5.8 | 2.5 | 0.8 | 3.3 | 0 | 0 | 0 | 4.2 | 13.3 | 0.8 |
| RUBBER | 13.3 | 4.2 | 0.0 | 91.7 | 24.2 | 0.8 | 0 | 0.0 | 0.0 | 0.0 |
| SILICON(IV) OXIDE | 0.0 | 0.0 | 0.0 | 0.0 | 0 | 0 | 0 | 0.0 | 0.0 | 10.0 |
| OTHER* | 9.2 | 1.7 | 0.0 | 372.5 | 23.3 | 28.3 | 8.3 | 1.3 | 2.1 | 28.8 |
| **TOTAL** | **42.5** | **9.2** | **1.6** | **600** | **124.9** | **63.3** | **14.9** | **6.8** | **17.1** | **91.8** |

*Depending on the sample includes: 1,4-butanediamine, neocryl xk-205, poly(1-butene), poly(neopentylene terephthalate), poly(styrene-ethylene-butylene), polyalkylbenzene, polycarbonate, polysorbate, polyvinylformal, PVA, polytetrafluoroethylene, and different mix of such polymers; please refer to supplementary data S1 for identifying polymers specific for each sample.

**Supplementary Table 2.** Average density of polymers identified in the follicular fluid and control water samples analysed.

| Polymer name | Average Density at 25ºC (g/mL) |
|---|---|
| 1,4-butanediamine | 0.877 |
| acrylonitrile butadiene styrene | 1.085 |
| alpex ck-450, cyclized rubber | 1.0 |
| cellulose acetate | 1.3 |
| cellulose propionate | 1.23 |
| color masterbatch polypropylene + 70% white pigment | 2 |
| cotton | 1.55 |
| duroftal vpi 2801/78 bac | 1.14 |
| dutral pmx 9705, copolymer epdm type | 0.87 |
| epichlorohydrin/ethylene oxide copolymer | 1.32 |
| hydroxybutyl methyl cellulose | 0.34 |
| keltan 314, copolymer epdm type | Not available |
| keltan 314, copolymer epdm type | 0.86 |
| methyl cellulose | 1.39 |
| neocryl xk-205 | 1.03 |
| neorez r 2020 | 1.03 |
| nylon 6,10 | 1.08 |
| olefin | 0.93 |
| pfa | 2.14 |
| poly(1-butene) | 0.95 |
| poly(acrylonitrile-co-butadiene-co-styrene) acrylonitrile 25% | 1.05 |
| poly(acrylonitrile-co-methyl acrylate) | Not available |
| poly(butylene terephthalate) | 1.3 |
| poly(ethylene terephthalate) | 1.385 |
| poly(ethylene-co-ethyl acrylate) 80:20 | 0.93 |
| poly(m-phenylene isophthalamide) | 1.38 |
| poly(styrene-ethylene-butylene) | 0.91 |
| poly(tetrafluoroethylene-co-perfluoro-(alkyl vinyl ether)) | 2.15 |
| poly(tetrafluoroethylene) | 2.0 |
| poly[(acrylic acid):(butyl acrylate):(methacryl amide):styrene] | 1.125 |
| poly[ethylene-co-(vinyl acetate)] | 0.98 |
| polyalkylbenzene | 0.853 |
| polyamide | 1.3 |
| polycarbonate | 1.2 |
| polyester | 1.38 |
| polyethylene | 0.91 |
| polyethylene glycol 200 | 1.124 |
| polyethylene terephthalate | 1.38 |
| polyethyleneimine | 1.07 |
| polyoxyethylene(20) sorbitan monopalmitate | 1.1 |
| polyoxyethylene(8) octylphenyl ether | 1.055 |
| polypropylene | 0.9 |
| polysorbate 80 standard | 1.06 |
| polystyrene | 1.05 |
| polyurethane | 1.7 |
| polyvinyl chloride | 1.38 |
| polyvinylformal | 1.23 |
| propanolamine | 0.9 |
| silicon(iv) oxide | 2.648 |
| styrene/acrylonitrile copolymer 75:25 | Not available |
| sympatex | 1.27 |
| synthalan ls 768 | 1.041 |
| teflon - polytetrafluoroethylene | 2.2 |
| viacryl sc 166 | 0.97 |
| vistalon 8510, copolymer epdm type | 1.19 |
